# Tract-explainable and underexplained synchrony play complementary roles in the functional organization of the brain

**DOI:** 10.64898/2026.03.19.712967

**Authors:** Jiebin Luo, Xin Zeng, Yue Xiong, Yuanhang Xu, Changsong Zhou, Yanjiang Wang, Dezhong Yao, Daqing Guo

## Abstract

Macroscale functional connectivity emerges from a complex interplay between structural wiring and non-tract-mediated mechanisms. However, the intrinsic entanglement of these contributions obscures their specific roles in brain organization. Here, we introduce a computational modeling framework to disentangle functional synchrony into two fundamentally distinct components: tract-explainable and tract-underexplained synchrony. Validated across two large-scale human cohorts (*n* = 1214) and an independent marmoset dataset (*n* = 24), this dissociation reveals an evolutionarily conserved architectural principle. We show that tract-explainable synchrony aligns closely with structural connectomes to facilitate global integration. Conversely, tract-underexplained synchrony drives local modularity and is anchored by multiscale cortical similarity, encompassing microstructural, receptor, and transcriptomic profiles. Crucially, these components gradually dissociate from sensori-motor to higher-order association cortices, with the importance of tract-underexplained synchrony progressively increasing. Furthermore, the tract-underexplained synchrony exhibits greater individual variability and demonstrates a stronger association with individual cognitive performance. Ultimately, tract-based and non-tract-mediated mechanisms serve distinct yet complementary roles, jointly shaping a functional organization that balances conserved macroscopic stability with higher-order cognitive flexibility.

## 1 Introduction

Functional connectivity (FC), which describes the statistical dependence of neural activity between brain regions, provides a powerful window into the intrinsic organization of large-scale brain networks [1–3]. The currently prevalent paradigm quantifies this connectivity using resting-state functional magnetic resonance imaging (rs-fMRI) by calculating Pearson correlation coefficients between regional brain activities [4, 5]. By revealing coordinated activity across distant cortical regions, these spatial patterns have been widely utilized to characterize cognitive architectures, individual variability, and clinical phenotypes [6–8]. However, because FC remains a purely statistical construct rather than a tangible entity, a fundamental question in systems neuroscience is how the brain leverages underlying physical interactions to generate these macroscopic connectivity patterns.

A classical and dominant paradigm in neuroscience posits that these functional patterns emerge from the coordinated dynamics of discrete, functionally specialized neuronal populations, which are interconnected by an intricate array of axonal fibers [9, 10]. In this view, the structural organization of the brain is the primary determinant of its functional repertoire. At the macroscale, this structural blueprint is commonly approximated by structural connectivity (SC) derived from diffusion magnetic resonance imaging (MRI), which estimates the density of white-matter fiber pathways linking cortical regions [11]. Consistent with the classical paradigm, a broad range of approaches centered on structural connectivity have demonstrated that SC shapes key features of FC, including its spatial modes and network topology [12–14].

Despite its explanatory power, a substantial structure–function mismatch persists. Mounting evidence indicates that this SC–FC divergence is far from random, displaying intrinsically organized patterns across cortical hierarchies, populations, and developmental stages [15–18]. Specifically, structural wiring tightly constrains functional coupling in unimodal sensorimotor cortices, but shows limited explanatory power in transmodal regions such as the default mode network [19]. To bridge this explanatory gap, recent studies have highlighted the profound influence of non-tract factors, demonstrating that cortical geometry and spatial gene expression profiles also fundamentally shape functional organization [20, 21]. Furthermore, integrating these non-structural features with anatomical connectivity significantly improves the characterization of empirical FC. For instance, predictive models achieve superior fits by incorporating inter-regional similarities in receptor distributions and microstructural profiles [22, 23]. Similarly, large-scale brain models have advanced our understanding of macroscale dynamics by parameterizing regional heterogeneity through receptor-mediated excitation-inhibition ratios or intracortical myelination [24, 25]. Together, these findings suggest that functional connectivity is shaped jointly by structural wiring and additional non-tract influences. However, because these influences are intrinsically intertwined in empirical FC, it remains difficult to disentangle their respective contributions to large-scale functional organization.

To address this challenge, we developed a large-scale brain modeling-based framework to decompose empirical functional synchrony into two complementary components. By incorporating tract-mediated interactions alongside regional background influences, the framework yields tract-explainable synchrony (Syn_TE_), which captures coupling constrained by structural wiring, and tract-underexplained synchrony (Syn_TU_), which isolates functional coordination less directly explained by anatomical tracts. We systematically validated this framework across two large human cohorts (total *n* = 1214) and a marmoset dataset (*n* = 24), demonstrating the high cross-species reproducibility of these components.

Our analyses reveal that Syn_TE_ and Syn_TU_ capture distinct yet complementary aspects of functional organization across the cortex. Specifically, while Syn_TE_ facilitates global integration, Syn_TU_ promotes regional specialization. Spatially, the two components gradually dissociate along the sensorimotor-association hierarchy, with Syn_TU_ becoming increasingly prominent. Crucially, while Syn_TE_ predominantly captures stable connectivity patterns shared across the population, Syn_TU_ demonstrates heightened sensitivity to individual-specific variability and cognitive performance. Together, these findings provide a new perspective on how large-scale brain function emerges from the interplay between a relatively fixed anatomical scaffold and flexible biological modulation, offering a mechanistic framework for understanding the diverse origins of functional connectivity.

## 2 Results

### 2.1 Model-Based Framework for Extracting Tract-Explainable and Underexplained Functional Synchrony

Our extraction framework builds on a large-scale brain modeling approach (see Methods for details). In this study, we employed the heterogeneous dynamic mean-field model (DMF) to simulate the local dynamics of each brain area [26]. Introducing regional heterogeneity improved the ability of large-scale brain model to capture region-specific functional characteristics and to more accurately reproduce empirical FC. To maintain computational tractability while preserving this heterogeneity, we adopted a low-dimensional parameterization informed by macroscale cortical gradients and myelin maps [26], and further linearized the model dynamics to enable efficient individual-level optimization and subsequent estimation of distinct synchrony components.

Based on the linearized model, we optimized the individual-level parameters to match simulated and empirical FC (Fig. 1A). Classical large-scale brain models typically assume that interregional functional synchrony is fully constrained by SC. However, at the macroscale, diffusion MRI–derived SC cannot fully account for empirical FC, due to both methodological limitations of tractography and additional non-axonal influences such as cortical proximity and homologous or symmetric projections. To incorporate these additional influences, we allowed the covariance matrix of the background stochastic noise (*v*), denoted *C_v_*, to be non-diagonal. Combining the estimated *C_v_* with the optimized model enabled us to derive two distinct functional synchrony matrices (see Methods): tract-explainable synchrony (Syn_TE_) and tract-underexplained synchrony (Syn_TU_). Syn_TE_ captures the component of functional synchrony primarily accounted for by tractography-derived coupling, whereas Syn_TU_ reflects synchrony that is less directly explained by tractography and likely reflects the aggregated influence of additional factors (Fig. 1B). In additional experiments, we further demonstrated that our proposed framework can be generalized to other large-scale models with different free parameters and local dynamics of each brain area (see Fig. S1A, B for details).

**Fig. 1.**
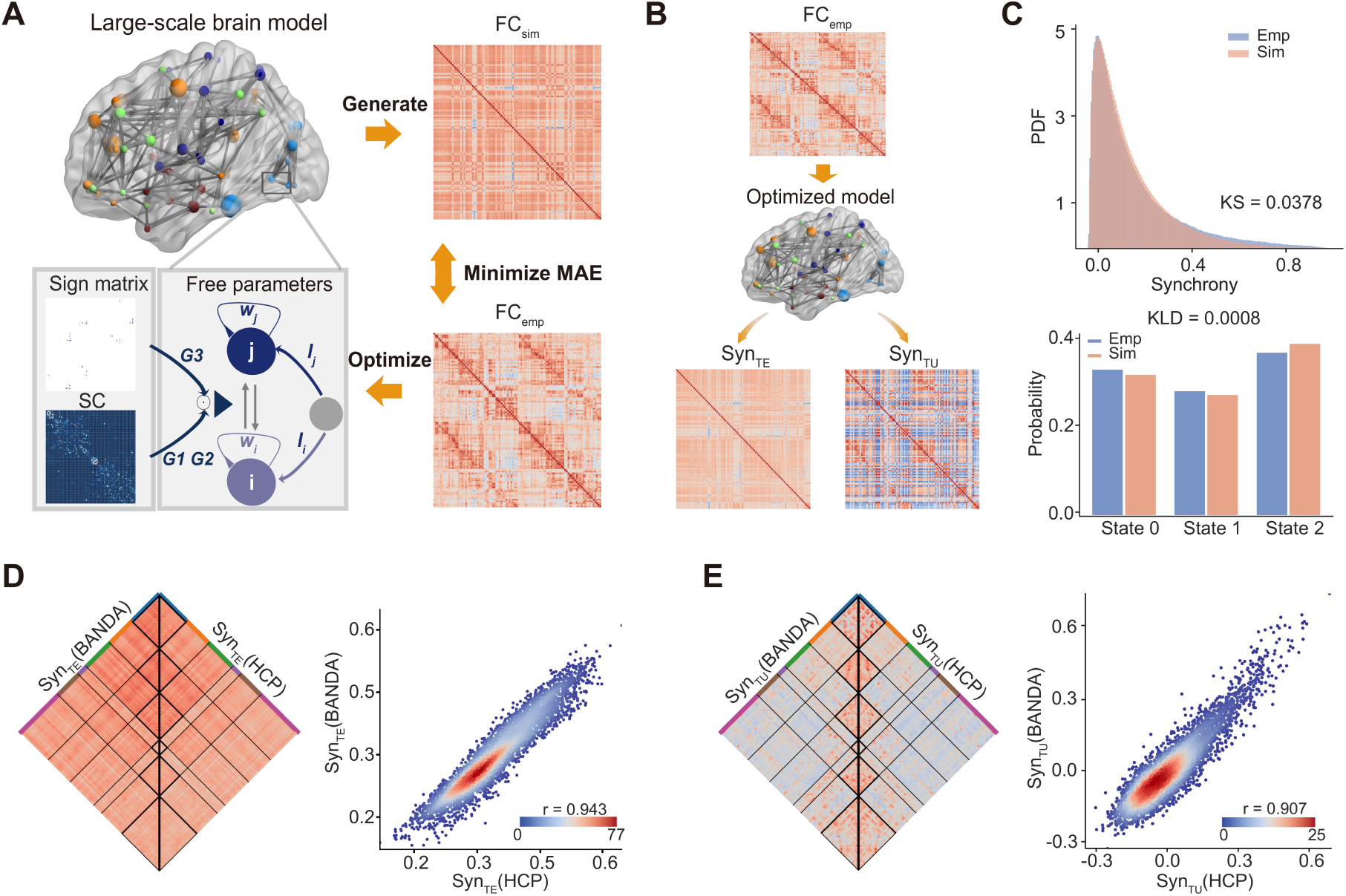
Model-based extraction of functional connectivity (FC) into tract-explainable and underexplained functional synchrony. **(A)** For each individual, the free parameters of a large-scale brain model with region-independent noise are optimized to minimize the mean absolute error (MAE) between simulated and empirical FC. **(B)** Using the optimized model, two complementary functional synchrony components are estimated from empirical FC: tract-explainable synchrony (Syn_TE_), reflecting inter-regional coupling primarily accounted for by diffusion MRI–derived tractography, and tract-underexplained synchrony (Syn_TU_), capturing synchrony not well explained by tractography and reflecting additional influences. Note that while empirical FC can be reconstructed from Syn_TE_ and Syn_TU_, empirical FC is not simply a linear sum of these two components. **(C)** Top: Probability density functions (pdf) of functional connectivity dynamics (FCD) from the individual-level model and empirical data are compared, with similarity quantified using the Kolmogorov–Smirnov (KS) statistic. Bottom: Probabilistic metastable substates (PMS) generated by the individual-level model are compared with empirical PMS, and their distributional similarity is quantified using the Kullback–Leibler divergence (KLD). **(D)** Left: Group-averaged Syn_TE_ from the Human Connectome Project (HCP) and Boston Adolescent Neuroimaging of Depression and Anxiety (BANDA) datasets. Right: Syn_TE_ estimates from the two datasets show high consistency (*r* = 0.943). **(E)** Left: Group-averaged Syn_TU_ from the HCP and BANDA datasets. Right: Syn_TU_ estimates from the two datasets show high consistency (*r* = 0.907).

To evaluate whether the two extracted functional synchrony matrices preserved the information contained in empirical FC, we simulated neural activity using these matrices. As shown in Fig. S1C, the simulated FC of a randomly selected participant closely matched the corresponding empirical FC. Importantly, although model optimization and estimation of *C_v_* relied solely on static empirical FC, the proposed framework also captured dynamic functional features, successfully reproducing functional connectivity dynamics (FCD) and probabilistic metastable substates (PMS) that closely resembled empirical observations (Fig. 1C). These results demonstrate that our model-based extraction framework effectively preserves individual-level brain functional characteristics, encompassing both static and dynamic aspects of functional connectivity.

We further assessed the cross-dataset consistency of Syn_TE_ and Syn_TU_. Group-level Syn_TE_ and Syn_TU_ were computed separately using the Human Connectome Project (HCP) dataset and the Boston Adolescent Neuroimaging of Depression and Anxiety (BANDA) dataset. The results revealed a high degree of cross-dataset consistency for both components, with strong correlations observed for Syn_TE_ (*r* = 0.943, *p <* 0.001) and Syn_TU_ (*r* = 0.907, *p <* 0.001) (Fig. 1D, E). These findings indicate that Syn_TE_ and Syn_TU_ capture robust and generalizable population-level functional synchrony patterns, and further support the reliability and stability of the proposed model-based extraction framework.

### 2.2 Distinct Biological Alignments of Tract-Explainable and Underexplained Functional Synchrony

From a model optimization perspective, Syn_TE_ should exhibit a stronger association with tractography-derived SC, because inter-regional coupling in the model is primarily shaped by SC. However, the optimization procedure does not impose a strict separation between tractography-related and other influences, meaning that Syn_TE_ may still contain non-tractography effects and Syn_TU_ may retain some SC dependence. We therefore tested whether the two components show differential associations with SC. At the group level, Syn_TE_ showed a stronger correlation with SC (*r* = 0.49, *p <* 0.001; Fig. 2A, top) than Syn_TU_ (*r* = 0.25, *p <* 0.001; Fig. 2A, middle). To assess robustness, the 1,000 HCP participants were randomly divided into 100 groups, and the same grouping scheme was used in all subsequent analyses. Because spatial proximity inflates inter-regional associations, Euclidean distance was regressed out in all analyses. Across groups, the association between Syn_TE_ and SC remained significantly stronger than that between Syn_TU_ and SC (*t*(99) = 23.983, *d* = 2.398, *p <* 0.001; Fig. 2A, bottom).

**Fig. 2.**
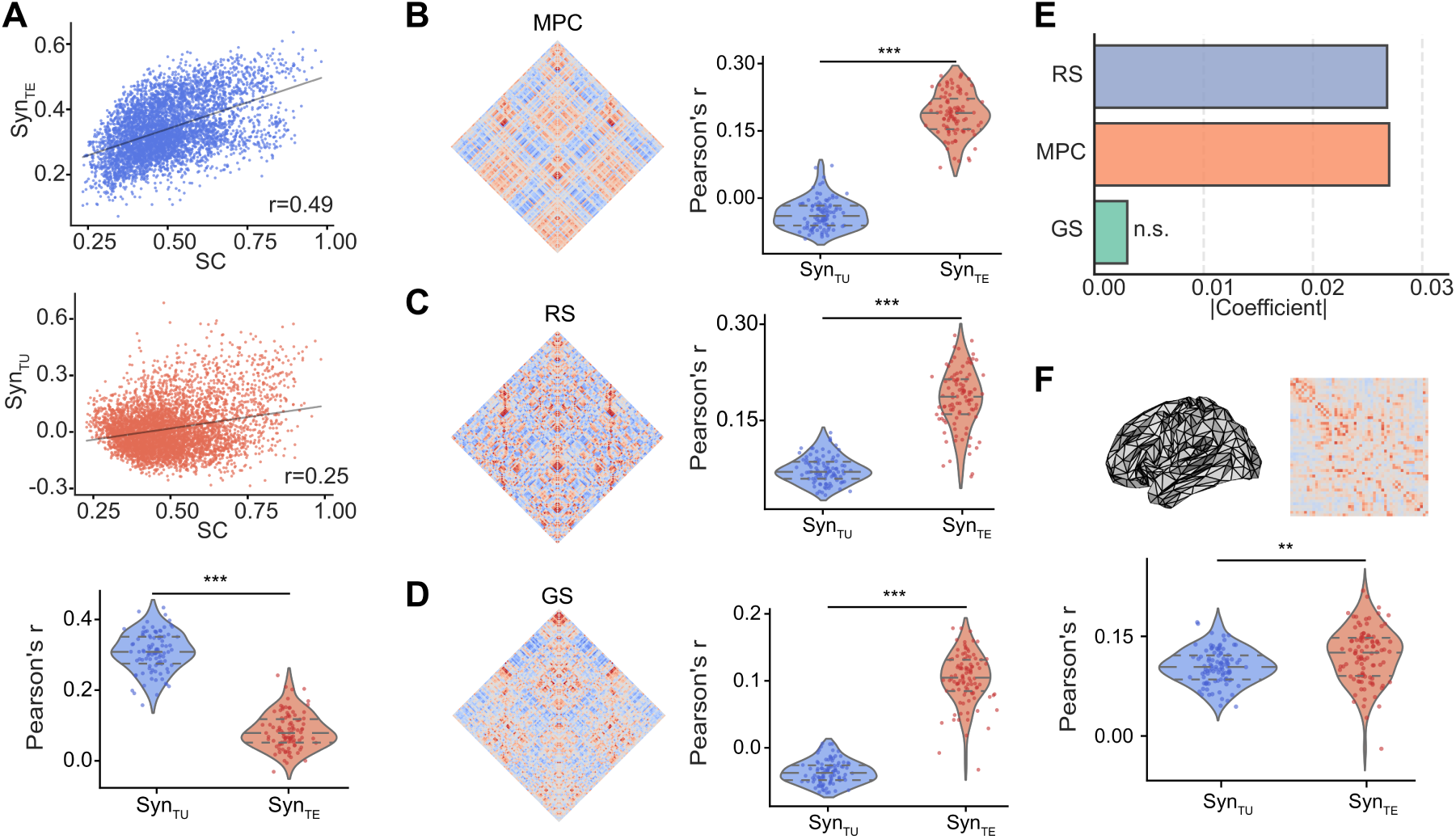
Differential associations of Syn_TE_ and Syn_TU_ with structural connectivity and multiscale cortical similarity features. **(A)** Top: Group-averaged correlation between Syn_TE_ and tractography-derived structural connectivity (SC) (*r* = 0.49, \*\*\**p <* 0.001). Middle: Corresponding correlation between Syn_TU_ and SC (*r* = 0.25, \*\*\**p <* 0.001). Bottom: After randomly splitting the HCP dataset into 100 groups and regressing out Euclidean distance, the association between Syn_TE_ and SC remained significantly stronger than that between Syn_TU_ and SC (paired-samples *t*-test, *t*(99) = 23.983, *d* = 2.398, \*\*\**p <* 0.001). The same distance-regression procedure was applied to all subsequent analyses. **(B)** Left: Microstructure profile covariance (MPC) matrix. Right: Syn_TE_ showed a significantly weaker correlation with MPC than Syn_TU_ (paired-samples two-tailed *t*-test, *t*(99) = *−*31.399, *d* = *−*3.140, \*\*\**p <* 0.001). **(C)** Left: Receptor similarity (RS) matrix. Right: The correlation between Syn_TE_ and RS was significantly weaker than that between Syn_TU_ and RS (paired-samples two-tailed *t*-test, *t*(99) = *−*21.307, *d* = *−*2.131, \*\*\**p <* 0.001). **(D)** Left: Gene-expression similarity (GS) matrix for the left hemisphere. Right: Syn_TE_ exhibited a significantly weaker correlation with GS than Syn_TU_ (paired-samples two-tailed *t*-test, *t*(99) = *−*16.442, *d* = *−*1.644, \*\*\**p <* 0.001). **(E)** Multilinear regression analysis showing that Syn_TU_ is strongly associated with MPC and RS, but not with GS. **(F)** Top: Cortical geometry and geometry-constrained FC generated by a neural field model using data and parameters from Pang et al. Bottom: Syn_TE_ showed a significantly weaker correlation with geometry-constrained FC than Syn_TU_ (paired-samples two-tailed *t*-test, *t*(99) = *−*3.472, *d* = *−*0.347, \*\**p <* 0.01).

To examine whether Syn_TU_ preferentially reflects non-tractography influences, we next compared the associations of Syn_TE_ and Syn_TU_ with several cortical similarity measures not directly captured by macroscale tractography: microstructure profile covariance (MPC), receptor similarity (RS), and gene-expression similarity (GS). Syn_TE_ showed a significantly weaker association with MPC than Syn_TU_ (*t*(99) = *−*31.399, *d* = *−*3.140, *p <* 0.001; Fig. 2B, right). A similar pattern was observed for RS (*t*(99) = *−*21.307, *d* = *−*2.131, *p <* 0.001; Fig. 2C, right) and GS (*t*(99) = *−*16.442, *d* = *−*1.644, *p <* 0.001; Fig. 2D, right). Multilinear regression further showed that Syn_TU_ was strongly associated with MPC and RS, whereas no significant association with GS was detected (Fig. 2E). Notably, these differential associations remained consistent even without regressing out the influence of interregional Euclidean distance (Fig. S2). Because FC is constrained by cortical geometry, we additionally examined whether Syn_TU_ aligns more strongly with geometry-driven synchrony. Using model parameters from Pang et al. [20], we generated FC constrained by cortical geometry using a neural field model. Syn_TE_ showed a significantly lower correlation with this geometry-constrained FC than Syn_TU_ (*t*(99) = *−*3.472, *d* = *−*0.347, *p <* 0.01; Fig. 2F, bottom).

Together, these results demonstrate a clear dissociation between Syn_TE_ and Syn_TU_. Specifically, Syn_TE_ is preferentially aligned with tractography-derived structural connectivity, whereas Syn_TU_ captures functional synchrony that is less directly explained by tractography and reflects the aggregated contribution of additional influences. These findings validate the effectiveness of the proposed framework in distinguishing tract explainable and underexplained components of large scale functional synchrony.

### 2.3 Topological and Gradient Dissociations of Syn_TE_ and Syn_TU_

As distinct component of functional synchrony extracted from empirical functional connectivity, Syn_TE_ and Syn_TU_ may capture complementary aspects of large-scale brain organization. To test this possibility, we first examined these synchrony component from a network perspective using graph-theoretical analysis. Considering the weighted and signed nature of functional brain networks, we adopted the modularity and connection diversity metrics proposed by Rubinov and Sporns as complementary metrics [27], because they capture the fundamental principles of brain organization: functional segregation and integration. Graph-theoretical analysis revealed a clear functional dissociation between the two components. Syn_TU_ showed significantly higher modularity compared to Syn_TE_ (*t*(99) = 82.06, *d* = 8.21, *p <* 0.001; Fig. 3A), indicating that Syn_TU_ preferentially supports functional specialization and segregation. Conversely, Syn_TE_ exhibited a significantly higher average connection diversity compared to Syn_TU_ (*t*(99) = 80.51, *d* = 8.05, *p <* 0.001; Fig. 3B), indicating that tract-explainable synchrony plays a dominant role in global communication and integration. These findings were consistently observed at the individual level (Fig. S3). To ensure the findings were not atlas-dependent, we replicated these results using a higher-resolution parcellation (Schaefer-400 atlas; Fig. S4).

**Fig. 3.**
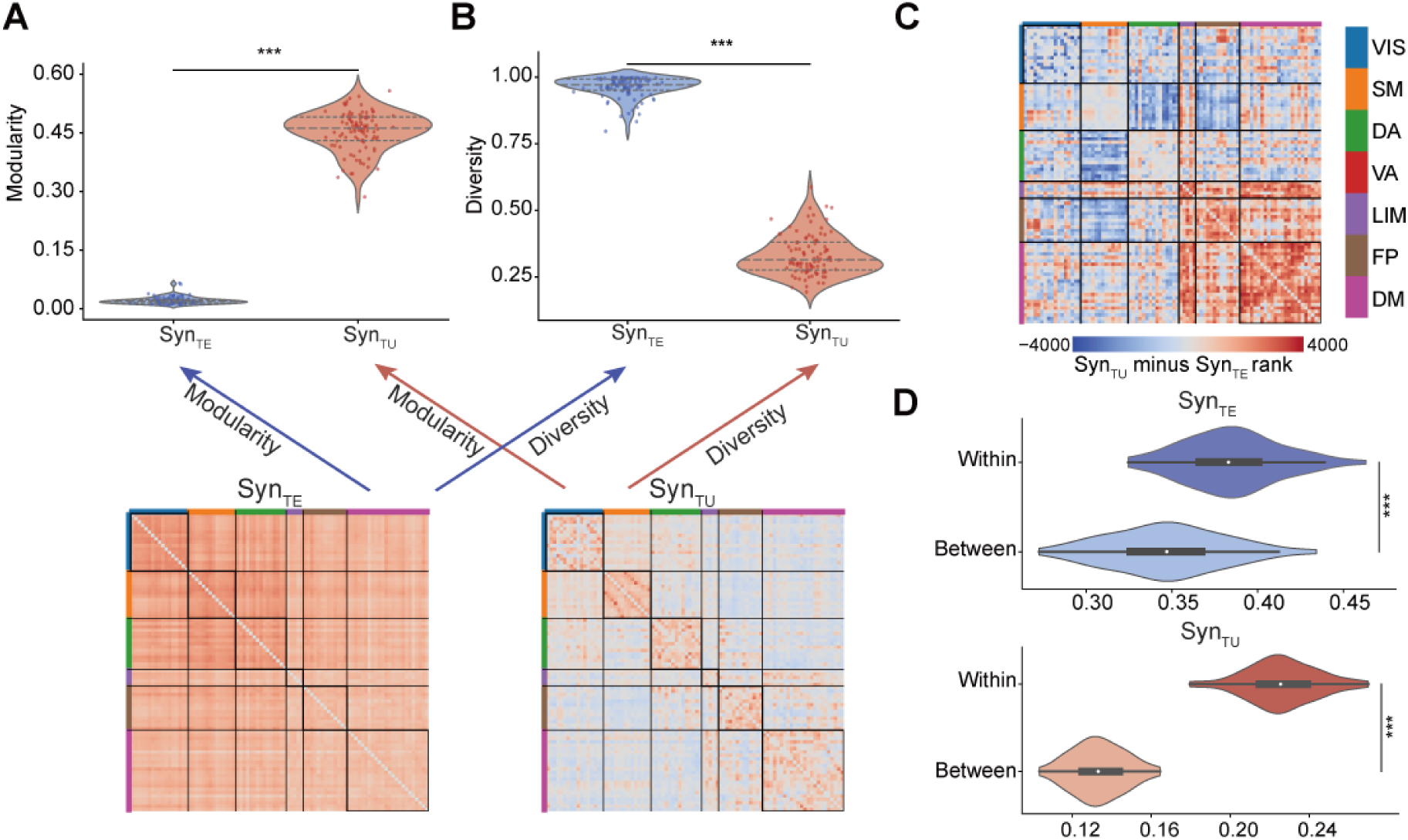
Distinct functional synchrony component exhibit dissociable network properties. **(A)** Syn_TU_ exhibited significantly higher modularity than Syn_TE_ (*t*(99) = 82.06, *d* = 8.21, \*\*\**p <* 0.001), indicating that tract-underexplained synchrony supports greater functional segregation. **(B)** Syn_TE_ exhibited a significantly higher average connectivity diversity than Syn_TU_ (*t*(99) = 80.51, *d* = 8.05, \*\*\**p <* 0.001), suggesting that tract-explainable synchrony plays a dominant role in global functional integration. **(C)** Matrix of Syn_TU_–Syn_TE_ gradient scores for each connection. The score was defined as the edge rank difference (Syn_TU_ rank minus Syn_TE_ rank) following the logic of Luppi et al.. Red (warm) colors indicate connections dominated by Syn_TU_, while blue (cool) colors indicate dominance of Syn_TE_. Regions are ordered by the Schaefer 7-network parcellation. **(D)** Top: Syn_TE_ showed significantly stronger within-resting-state network (RSN) synchrony than between-RSN synchrony (*t*(99) = 19.63, *d* = 1.96, \*\*\**p <* 0.001). Bottom: Syn_TU_ likewise exhibited significantly stronger within-RSN synchrony compared with between-RSN synchrony (*t*(99) = 65.63, *d* = 6.56, \*\*\**p <* 0.001).

Beyond these global topological differences, the relative contributions of tract-explainable and underexplained synchrony varied substantially across individual connections. To capture this heterogeneity and quantify the relative dominance of each synchrony mode, we computed a connection-wise Syn_TU_–Syn_TE_ gradient metric. This metric, representing an edge rank gradient calculated following the logic of Luppi et al. [28], was defined as the difference between the rank of a given edge in the Syn_TU_ matrix and its rank in the Syn_TE_ matrix (see Methods). This connection-wise gradient revealed a clear and spatially organized network-level structure (Fig. 3C), with the relative contribution of Syn_TU_ progressively increasing across functional systems, consistent with a shift from lower-order to higher-order network organization. Together, these findings indicate that tract-based and non-tract-based influences differentially shape functional interactions across the whole-brain connectome.

Despite these pronounced differences in global topology and connection-wise contributions, both synchrony component exhibited significantly stronger within-network than between-network coupling (Fig. 3D), consistent with canonical large-scale network organization [29]. This result indicates that although Syn_TE_ and Syn_TU_ differ in their topological profiles and spatial distributions, they both preserve the fundamental principle of preferential intra-network connectivity that characterizes human brain functional architecture.

Having established these fundamental network properties, we next examined the functional embedding of Syn_TE_ and Syn_TU_ using cortical gradients to understand how these synchrony component map onto established functional hierarchies. In the low-dimensional manifold defined by the first two gradients, Syn_TU_ exhibited pronounced within-network clustering and between-network separation (Fig. 4A), closely resembling the organization observed in empirical FC [3]. In contrast, the embedding of Syn_TE_ appeared more distributed across the manifold; while vis subnetwork maintained clear separation from other systems, the remaining networks showed higher degrees of spatial overlap. Accompanying these distinct clustering patterns, their principal gradients diverged markedly: the Syn_TE_ gradient extended from somatomotor to visual regions, whereas the Syn_TU_ gradient extended from somatomotor to default-mode regions. To interpret these differences, we projected NeuroSynth-derived cognitive and behavioral terms onto the principal gradients. Motor, action, and pain-related terms occupied similar positions at one end of both gradients, whereas the opposite ends diverged: the terminal end of the Syn_TE_ gradient aligned with visual-related regions, while the terminal end of the Syn_TU_ gradient was enriched for regions associated with higher-order cognitive and memory functions (Fig. 4B).

**Fig. 4.**
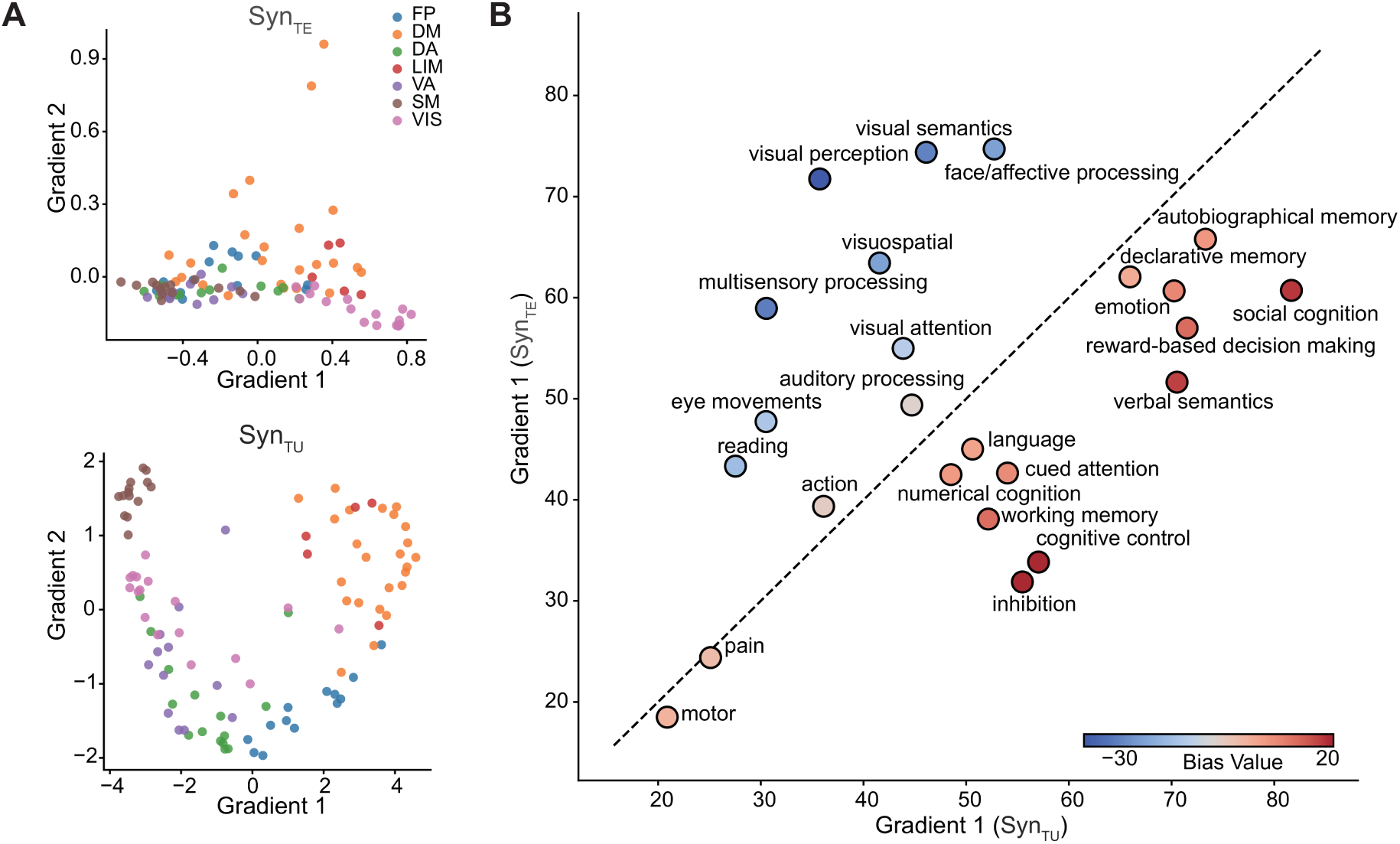
Gradient-level dissociation between Syn_TE_ and Syn_TU_. **(A)** Top: The first and second functional gradients derived from Syn_TE_. In this low-dimensional manifold, the visual sub-network maintains clear separation while other systems exhibit higher degrees of spatial overlap. Bottom: The first and second functional gradients derived from Syn_TU_, showing pronounced within-network clustering and between-network separation. **(B)** Centroid positions of cognitive and behavioral terms from NeuroSynth projected onto the principal gradients of Syn_TE_ and Syn_TU_. The color of each point indicates the positional deviation of the corresponding term between the two gradient spaces, highlighting domain-specific differences in gradient organization between Syn_TE_ and Syn_TU_.

### 2.4 The Syn_TE_-Syn_TU_ Axis Aligns with Hierarchy and Synaptic Organization

Syn_TE_ and Syn_TU_ reflect the influences of distinct factors on brain functional connectivity and reveal complementary aspects of large-scale functional organization. Their dissociable functional gradients further suggest that cortical regions with different structural substrates and functional demands may rely differentially on these two synchrony component. To explicitly characterize how Syn_TE_ and Syn_TU_ relate to one another at the regional level, we quantified their relationship from two complementary perspectives: within-region spatial consistency and relative importance.

Within-region spatial consistency was measured using Explained–Underexplained Synchrony Consistency (EUSC), defined as the regional Spearman correlation between the connectivity profiles of Syn_TE_ and Syn_TU_. Relative importance was quantified using the Underexplained–Minus–Explained Rank (UMER), defined as the difference between the nodal strength rank in Syn_TU_ and that in Syn_TE_ (see Methods for detailed definitions). Theoretically, these two metrics capture distinct dimensions of functional organization. To verify that their observed close relationship reflects specific biological organization rather than mathematical redundancy, we compared the empirical results against a null model. We generated 1,000 surrogate networks by randomly shuffling the empirical connectivity matrices while preserving the degree distribution. Crucially, we found that the spatial correlation between EUSC and UMER maps in the null networks was substantially lower than that observed in the empirical data (*p <* 0.001; Fig. S5). This confirms that the strong coupling between spatial decoupling and functional dominance shifts is a non-trivial feature unique to the brain’s functional organization, rather than a mathematical artifact. Given this empirically robust convergence, we performed principal component analysis (PCA) on EUSC and UMER to parsimoniously capture their shared variance. The resulting Syn_TE_–Syn_TU_ Interaction Gradient, defined as the first principal component, explained 87% of the total variance. This gradient captures a systematic shift in brain organization: as one moves along the gradient, the spatial consistency between Syn_TE_ and Syn_TU_ progressively declines, while the relative importance of Syn_TU_ systematically increases (Fig. 5A).

**Fig. 5.**
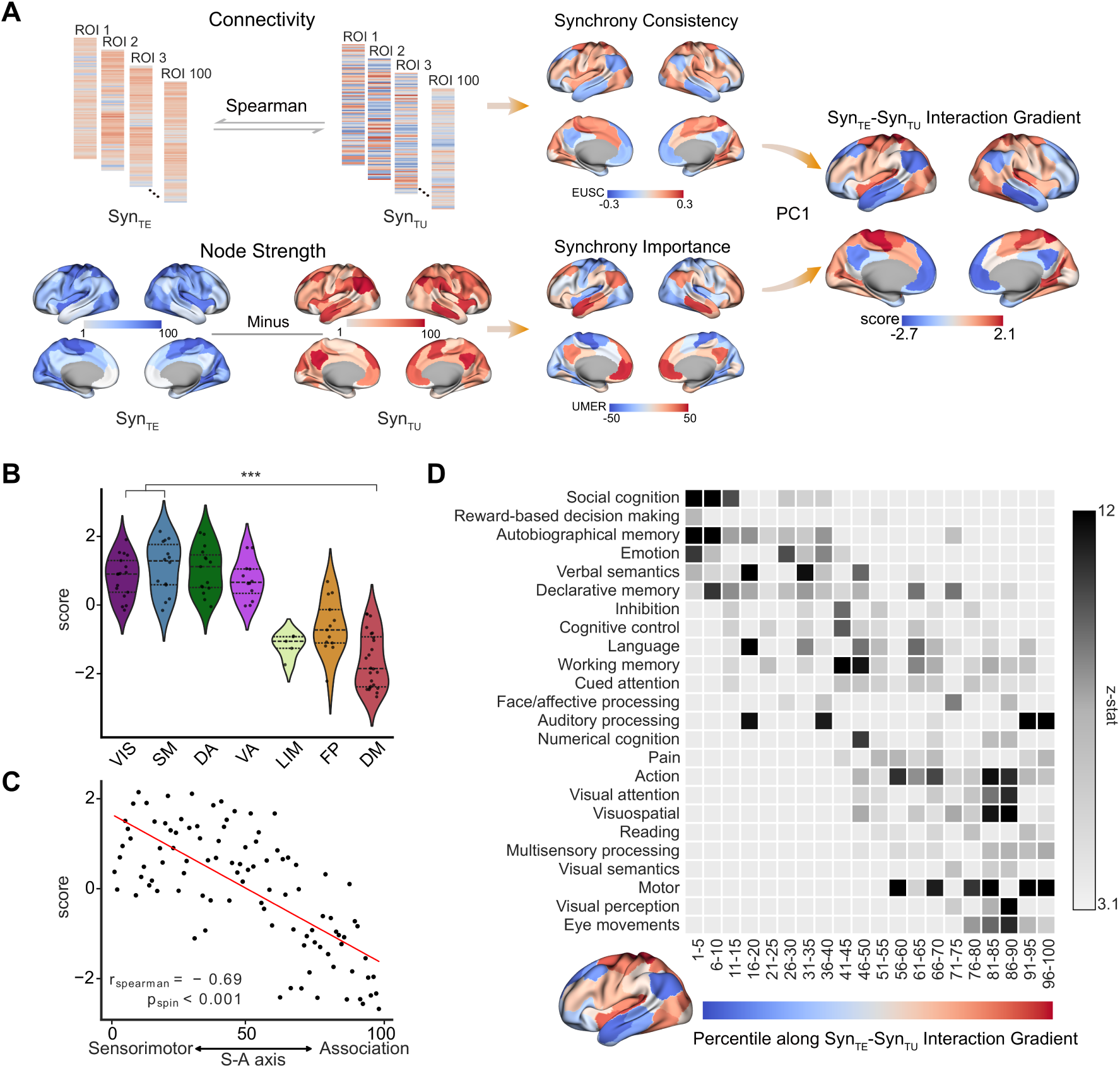
Cortical gradient capturing the interaction between Syn_TE_ and Syn_TU_. **(A)** The Syn_TE_–Syn_TU_ Interaction Gradient was defined as the first principal component derived from two complementary metrics: Explained–Underexplained Synchrony Consistency, which quantifies the regional Spearman correlation between Syn_TE_ and Syn_TU_, and the Underexplained–Minus–Explained Rank, representing the difference in nodal strength rank between the two synchrony modes. This dominant gradient explains 87% of the total variance, capturing a systematic transition where the consistency between Syn_TE_ and Syn_TU_ progressively decreases as the relative importance of Syn_TU_ increases. **(B)** The distribution of the Syn_TE_–Syn_TU_ Interaction Gradient scores across the seven canonical resting-state networks. The default mode (DM) network exhibits significantly lower gra-dient scores compared to the visual (VIS) and somatomotor (SM) networks (*p <* 0.001), signifying a more pronounced dissociation between Syn_TE_ and Syn_TU_ and a dominant relative contribution of Syn_TU_ within the DM network. **(C)** The Syn_TE_–Syn_TU_ Interaction Gradient is significantly correlated with the sensorimotor–association (S–A) cortical axis. **(D)** NeuroSynth-based meta-analysis of the gradient. Brain regions are ranked by their Syn_TE_–Syn_TU_ Interaction Gradient scores and partitioned into 20 equally sized bins (*x*-axis), while the *y*-axis presents cognitive and behavioral terms. Heatmap intensity reflects the mean task-activation *z*-scores within each bin, revealing a transition from basic sensorimotor functions to higher-order cognitive domains.

To assess the functional significance of this cortical gradient, we examined the topographic variation of the Syn_TE_–Syn_TU_ Interaction Gradient across canonical functional networks, cortical hierarchies, and cognitive domains. At the network level, the default mode (DM) network—commonly regarded as a higher-order association network—exhibited significantly lower gradient scores than the visual (VIS) and somatomotor (SM) networks (Fig. 5B), indicating greater dissociation between Syn_TE_ and Syn_TU_ and a stronger relative contribution of Syn_TU_ within association cortex. At the regional level, gradient scores showed a strong negative correlation with the position of cortical regions along the sensorimotor–association (S–A) axis (*r*_spearman_ = *−*0.69*, p*_spin_ *<* 0.001, two-tailed; Fig. 5C). Sensorimotor regions displayed higher Syn_TE_–Syn_TU_ consistency and greater reliance on Syn_TE_, whereas association regions exhibited increased dissociation and a progressively greater reliance on Syn_TU_. Consistent with this hierarchical organization, a NeuroSynth-based meta-analysis using the same 24 cognitive topic terms as in Margulies et al. revealed a systematic functional dissociation along the gradient [3]. Lower gradient scores were associated with higher-order cognitive domains, including social cognition, emotion, reward processing, and decision making, whereas higher gradient scores were linked to basic sensorimotor functions such as eye movements, motor control, and visual perception (Fig. 5D). Notably, these hierarchical associations were further validated using a more granular parcellation scheme (Schaefer-400 atlas), yielding consistent results (Fig. S6).

To elucidate the molecular and synaptic substrates underlying this cortical axis, we conducted a comprehensive gene set enrichment analysis. Strikingly, the most significantly enriched terms across multiple databases were overwhelmingly dominated by neuronal and synapse-related functions, including neuron projection, chemical synaptic transmission, and nervous system development (Fig. 6A). Crucially, the genes driving these top pathways universally exhibited robust negative correlations with the gradient scores. Building on these findings, we further examined whether the gradient was associated with distinct *in vivo* and *ex vivo* markers of synaptic organization. We considered three complementary metrics: synaptic density estimated using positron emission tomography (PET) with the synapse-specific radioligand [^11^C]UCB-J, synaptic development indexed by the cortical distribution of the glycolytic index (GI), and receptor characteristics captured by the principal gradient of neurotransmitter receptor distribution. Multiple linear regression revealed that synaptic development and the receptor principal gradient were independent predictors of the interaction gradient, whereas synaptic density was not (Fig. 6B). Given the established link between receptor gradients and receptor diversity, we further incorporated *ex vivo* autoradiographic receptor data. Receptor diversity, quantified as receptor entropy following Goulas et al. [30], showed a significant negative correlation with gradient scores (*r*_spearman_ = *−*0.48*, p <* 0.001, two-tailed; Fig. 6C), further supporting a close association between the Syn_TE_–Syn_TU_ axis and synaptic and receptor organization.

**Fig. 6.**
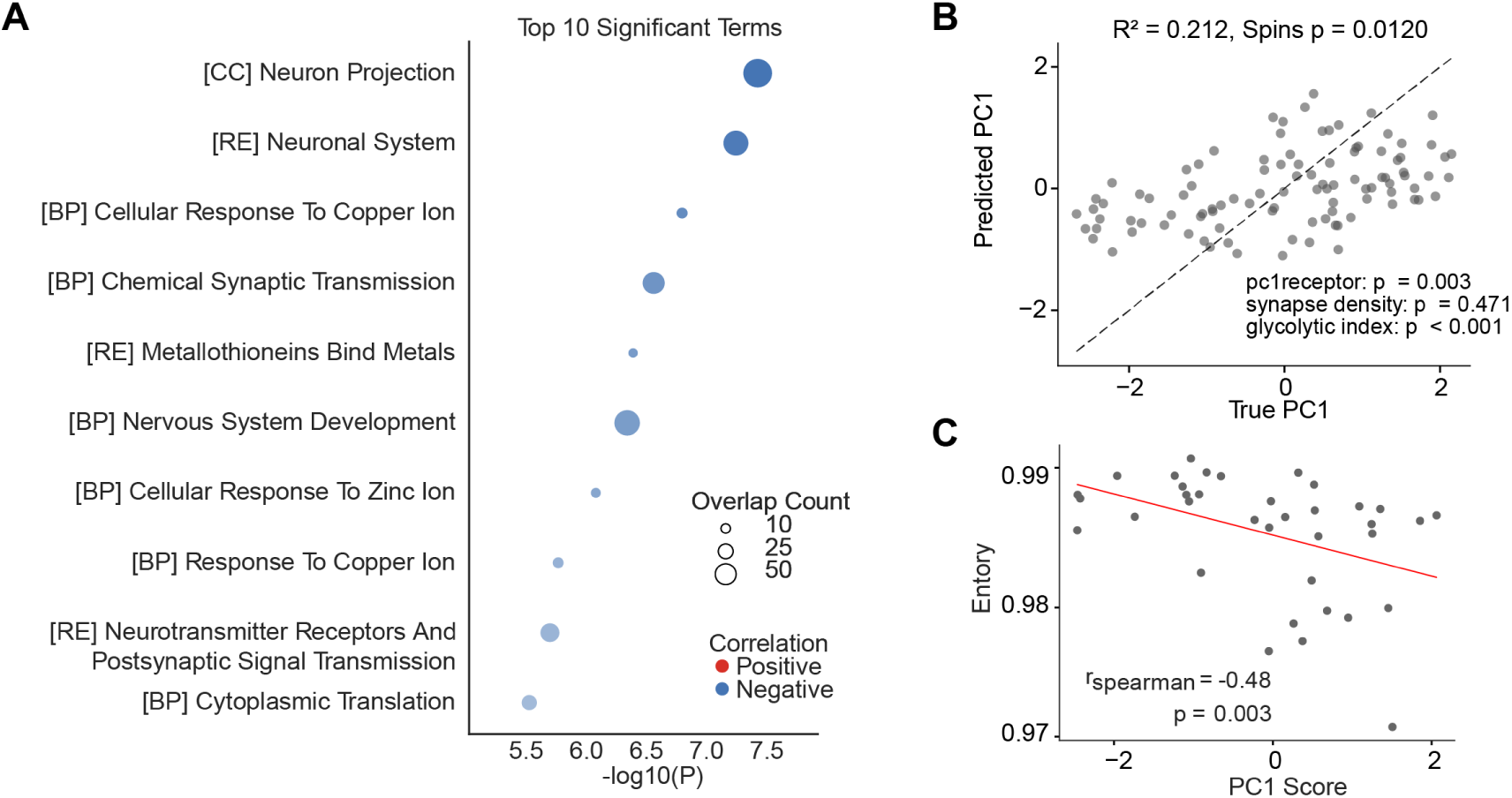
The Syn_TE_–Syn_TU_ Interaction Gradient is closely associated with synaptic organization. **(A)** Comprehensive gene set enrichment analysis associated with the Syn_TE_–Syn_TU_ interaction gradient. To capture a broader biological context, the analysis integrates multiple databases: Gene Ontology ([GO]), Biological Processes ([BP]), Molecular Functions ([MF]), Cellular Components ([CC]), and Reactome pathways ([RE]). The top ten significantly enriched terms across all combined datasets are shown. Bubble color indicates the direction of correlation (red = positive; blue = negative), and bubble size reflects the number of overlapping genes contributing to each term. **(B)** To further probe the synaptic basis of this gradient *in vivo*, multiple linear regression was performed using complementary synaptic metrics, including the principal gradient of neurotransmitter receptor distribution (*pc*1_receptor_), synaptic density estimated with the PET radioligand [^11^C]UCB-J, and the glycolytic index reflecting synaptic development. The model explained a significant proportion of variance in the Syn_TE_–Syn_TU_ interaction gradient, with pc1receptor and glycolytic index emerging as independent predictors, whereas synaptic density was not significant. **(C)** Consistent with the involvement of receptor organization, the Syn_TE_–Syn_TU_ interaction gradient showed a significant negative Spearman correlation with neurotransmitter receptor diversity (entropy) measured using *in vitro* quantitative autoradiography, providing independent *ex vivo* support for the association between the gradient and synaptic receptor architecture.

We further found that the Syn_TE_–Syn_TU_ Interaction Gradient is significantly correlated with the level of cortical developmental expansion (*r* = *−*0.40, *p*_spin_ = 0.021; Fig. S8A). Moreover, this gradient is closely associated with the heterogeneity of cortical temporal dynamics, as evidenced by significant correlations with intrinsic neural timescales (*r* = *−*0.50, *p*_spin_ = 0.012; Fig. S8B), a proxy for the duration of local temporal integration windows, and regional timescores (*r* = 0.43, *p*_spin_ = 0.027; Fig. S8C), a composite metric capturing the dominant gradient of MEG-derived dynamic features. Finally, the interaction gradient was linked to the mean power of specific frequency bands, including significant associations with delta and high-gamma power (Fig. S8D).

### 2.5 Tract-Underexplained Synchrony Reflects Individual Variability and Cognitive Performance

Given the pronounced group-level differences between Syn_TE_ and Syn_TU_, we next examined whether these two synchrony component capture distinct sources of inter-individual variability. Inter-individual similarity was quantified separately for Syn_TE_ and Syn_TU_ by computing pairwise correlations between the upper-triangular elements of individual connectivity matrices. We found that the inter-individual similarity of Syn_TE_ was substantially higher than that of Syn_TU_ (*t*(511, 565) = 1362.40, *d* = 1.91, *p <* 0.001; Fig. 7A). This high consistency of Syn_TE_ parallels the high inter-individual similarity observed in the empirical SC (Fig. S9), confirming that Syn_TE_ faithfully preserves the stable anatomical backbone shared across individuals. In contrast, Syn_TU_ captures greater inter-individual heterogeneity, suggesting it reflects functional idiosyncrasies that are not strictly constrained by white matter tracts.

**Fig. 7.**
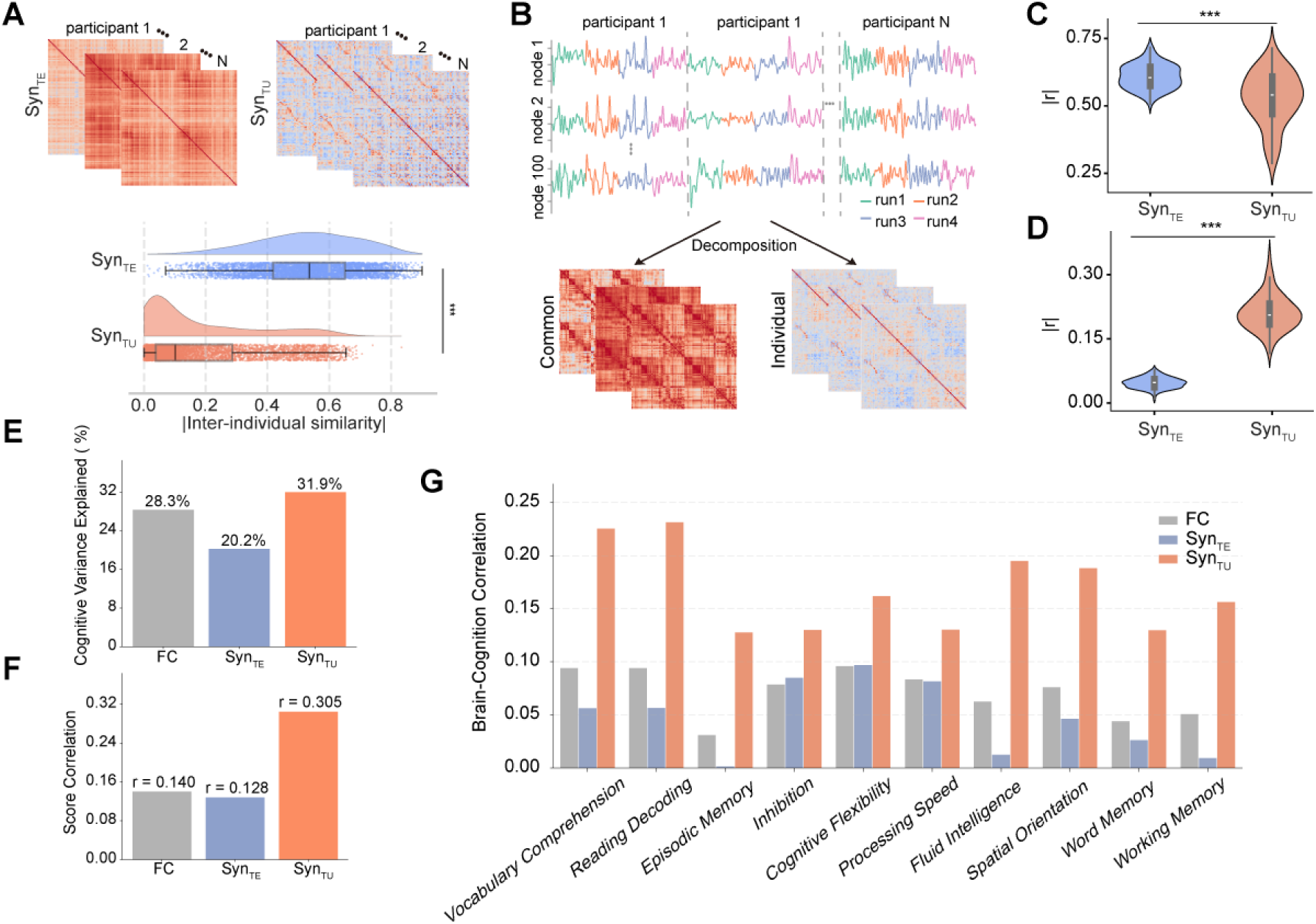
Individual-specific functional connectivity encoded by Syn_TU_ drives superior brain-cognition coupling. **(A)** Top: Connectivity matrices showing Syn_TE_ (left) and Syn_TU_ (right) estimated for representative individuals. Bottom: Quantitative comparison reveals that inter-individual similarity is substantially higher for Syn_TE_ than for Syn_TU_ (paired-samples two-tailed *t*-test, *t*(511, 565) = 1362.40, *d* = 1.91, \*\*\**p <* 0.001). **(B)** Schematic of the functional decomposition framework. Time series from four resting-state scans were concatenated across all participants, and the first six principal components were extracted to derive group-level shared co-activation patterns. Shared FC was computed within this low-dimensional subspace, whereas the residuals were used to estimate individual-specific FC. **(C)** Syn_TE_ exhibits a significantly stronger correlation with the shared FC component compared to Syn_TU_ (paired-samples two-tailed *t*-test, *t*(99) = 6.19, *d* = 0.62, \*\*\**p <* 0.001). **(D)** Conversely, Syn_TU_ shows a significantly stronger correlation with the individual-specific FC component compared to Syn_TE_ (paired-samples two-tailed *t*-test, *t*(99) = *−*38.51, *d* = *−*3.85, \*\*\**p <* 0.001). **(E)** Total cognitive variance explained by the first latent variable (LV1) derived from Partial Least Squares (PLS) analysis. **(F)** Brain-cognition score correlations (Pearson’s *r*) for LV1. **(G)** Brain-cognition correlation profiles across ten individual cognitive tasks. Bar charts represent the absolute correlation strength (*|r|*) between LV1 brain network scores and specific cognitive measures for FC, Syn_TE_, and Syn_TU_.

To directly test whether this dissociation reflects shared versus individual-specific functional components, we adopted the connectome caricatures framework, in which empirical BOLD time series are separated into a group-level common space and an individual-specific space (Fig. 7B; see Methods) [31]. This procedure yields a shared FC component and an individual-specific FC component unique to each participant. Syn_TE_ showed significantly stronger correspondence with the shared FC component than did Syn_TU_ (*t*(99) = 6.19, *d* = 0.62, *p <* 0.001; Fig. 7C). In contrast, Syn_TU_ exhibited substantially stronger correspondence with the individual-specific FC component compared with Syn_TE_ (*t*(99) = *−*38.51, *d* = *−*3.85, *p <* 0.001; Fig. 7D). Together, these convergent results establish a clear functional dissociation: Syn_TE_ preferentially preserves inter-individual shared components of functional connectivity, whereas Syn_TU_ emphasizes individual-specific functional information.

If Syn_TU_ indeed captures individual-specific aspects of functional organization, it should be particularly informative for explaining inter-individual differences in cognition. To test this hypothesis, we selected 10 tasks encompassing diverse domains of individual cognitive abilities [8] (Table S1). We employed a standard Partial Least Squares (PLS) analysis to identify linear combinations of connectivity features and cognitive scores that maximize their cross-covariance. The statistical significance of each latent variable (LV) was assessed using permutation testing. As shown in Fig. S10, the first latent variables for Syn_TU_, Syn_TE_, and FC were all statistically significant and accounted for the largest proportion of cross-covariance.

Interestingly, although LV1 of Syn_TU_ captured a smaller fraction of the overall cross-covariance compared to Syn_TE_ and FC, it explained a substantially higher proportion of the variance in cognitive scores (Fig. 7E). Concurrently, Syn_TU_ achieved a stronger brain-cognition score correlation for LV1 than both Syn_TE_ and FC (Fig. 7F). This discrepancy suggests that traditional FC and Syn_TE_ are likely dominated by macroscopic global patterns that are largely irrelevant to individual cognitive differences. Furthermore, directly evaluating the correlation between the LV1 brain network scores and individual cognitive measures revealed that Syn_TU_ exhibits consistently stronger correlation profiles across all ten cognitive tasks compared to both Syn_TE_ and FC (Fig. 7G). Ultimately, these findings demonstrate that by stripping away the anatomically constrained, group-shared backbone, the remaining tract-underexplained synchrony effectively captures individual-specific functional information that is highly relevant to human cognition.

### 2.6 Cross-Species Validation in the Marmoset

To explicitly evaluate whether the functional dissociation between Syn_TE_ and Syn_TU_ represents a fundamental principle of primate brain organization, we extended our analytical framework to an independent marmoset dataset [32]. Our results revealed that the topological features of these two synchrony components in the marmoset brain are highly concordant with our findings in humans. Specifically, Syn_TE_ exhibited significantly lower modularity compared to Syn_TU_ (*t*(23) = *−*8.45, *d* = *−*1.72, *p <* 0.001; Fig. 8A). Conversely, connection diversity was significantly higher in Syn_TE_ than in Syn_TU_ (*t*(23) = 9.16, *d* = 1.87, *p <* 0.001; Fig. 8B).

**Fig. 8.**
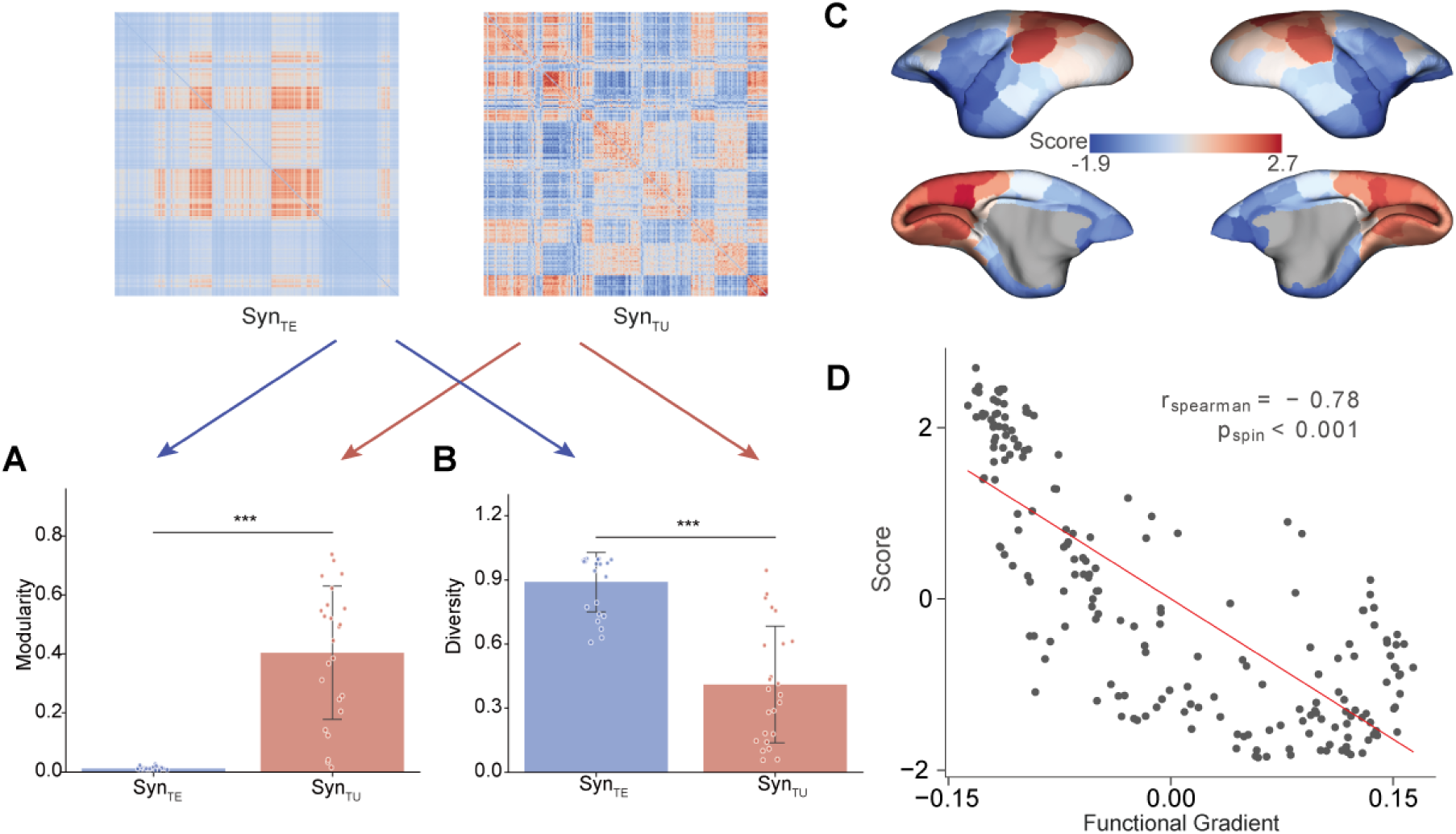
Topological properties and interaction gradients in the marmoset brain. Results demonstrate the cross-species conservation of the functional dissociation between Syn_TE_ and Syn_TU_ in the marmoset (*n* = 24). **(A)** The modularity of Syn_TE_ was significantly lower than that of Syn_TU_ (paired-samples *t*-test, *t*(23) = *−*8.45, *d* = *−*1.72, \*\*\**p <* 0.001). **(B)** The connection diversity (measured via participation coefficient) of Syn_TE_ was significantly higher than that of Syn_TU_ (paired-samples *t*-test, *t*(23) = 9.16, *d* = 1.87, \*\*\**p <* 0.001). **(C)** Cortical visualization of the Syn_TE_–Syn_TU_ interaction gradient across the marmoset cortex. **(D)** Spatial correlation between the Syn_TE_–Syn_TU_ interaction gradient and the intrinsic functional hierarchy (principal gradient) of the marmoset brain.

Beyond regional topology, we further computed the Syn_TE_–Syn_TU_ interaction gradient in marmosets and mapped it onto the cortical surface (Fig. 8C). Remarkably, spatial correlation analysis demonstrated that this interaction gradient is highly correlated with the principal functional gradient of the marmoset brain (*r*_spearman_ = *−*0.78, *p <* 0.001, two-tailed; Fig. 8D). Together, these robust cross-species replications confirm that the hierarchical and topological divergence between Syn_TE_ and Syn_TU_ is an evolutionarily conserved architectural feature across the primate lineage.

## 3 Discussion

In this study, we developed a large-scale brain modeling framework that decomposes empirical functional connectivity into two complementary components: tract-explainable synchrony (Syn_TE_) and tract-underexplained synchrony (Syn_TU_), thereby disentangling tract-mediated and non-tract-mediated contributions. By explicitly parameterizing non-tract influences through a non-diagonal background noise covariance structure, this approach addresses a fundamental limitation of functional connectivity, namely that it conflates multiple sources of influence and obscures their distinct roles. Our findings point to a core organizational principle of the human brain: a division between structurally constrained and structurally flexible interactions. Syn_TE_ reflects a conserved anatomical backbone that is highly consistent across individuals, supporting stable and globally integrative communication, whereas Syn_TU_ captures interactions that are less constrained by white-matter architecture and instead contribute to regional specialization, inter-individual variability, and cognitive relevance. Together, these results suggest that large-scale brain function emerges from the interplay between these two complementary systems, with structurally constrained interactions providing a stable, shared scaffold and non-tract-mediated inter-actions enabling functional diversity and individual-specific modulation, a principle further supported by cross-species validation.

The dissociation between Syn_TE_ and Syn_TU_ in network topology and physiological alignment indicates that macroscale functional interactions are unlikely to be governed by a single mode of physical communication. Instead, they appear to emerge from a multi-modal communication landscape in which tract-based and non-tract-based influences coexist. This principle is conceptually consistent with the well-established coexistence of wiring transmission (WT) and volume transmission (VT) at the neurobiological level [33]. In this interpretive framework, Syn_TE_ reflects a wiring-based mode of interaction rigidly constrained by diffusion MRI–derived structural connectivity, supporting stable global integration through efficient point-to-point pathways. In contrast, Syn_TU_ captures a form of functional coordination that is less directly constrained by anatomical tracts and more closely aligned with regional molecular and microstructural properties. We propose that this coordination likely arises from shared physiological inputs acting as natural sources of co-variance, including unmodeled subcortical projections (e.g., ascending neuromodulatory systems) and genuine VT [34–36]. Importantly, VT is not restricted to local diffusion; it encompasses a spectrum of non-synaptic communication ranging from short-range diffusion within the extracellular fluid to long-range transmission mediated by the cerebrospinal fluid (CSF) [37]. Crucially, the strong associations between Syn_TU_ and microstructure profile covariance, receptor similarity, and gene-expression similarity suggest that this coordination is facilitated by molecular homophily. Whether driven by widespread subcortical axons or fluid-mediated chemical signals, regions with similar cytoarchitecture or receptor profiles may exhibit convergent physiological responses even in the absence of direct structural connections [21, 38]. Thus, Syn_TU_ reflects a distal coordination mechanism shaped by molecular homophily, complementing the tract-based interactions captured by Syn_TE_. Together, these findings support a view of large-scale brain communication in which fiber-based conduction and non-tract-mediated influences represent complementary dimensions of functional organization.

The regional variation in the relationship between Syn_TE_ and Syn_TU_ is closely tied to the cortical hierarchy, revealing a hierarchical logic in large-scale communication across the cerebral cortex. At the lower end of this hierarchy, particularly in primary sensorimotor regions, the two synchrony components exhibit greater concordance. This alignment likely arises because, in these unimodal areas, non-tract-mediated influences do not operate independently but rather spatially overlap with and reinforce the point-to-point structural pathways. Such a synergistic configuration—where modulatory gradients are co-extensive with axonal bundles—is well suited for rapid, stimulus-driven processing, ensuring that chemical environments amplify immediate neural responses to external input with high fidelity [28, 39]. As the hierarchy ascends toward higher-order association regions, Syn_TE_ and Syn_TU_ progressively decouple, and Syn_TU_ becomes relatively more prominent. Unlike the rigid reinforcement required in sensory cortex, higher-order processing demands flexibility [40]. In these regions, functional coordination must transcend fixed anatomical pathways to support abstract cognition. By decoupling from the structural backbone, Syn_TU_ facilitates the emergence of diverse functional configurations that are not strictly dictated by hard-wiring [41]. Although our model does not explicitly simulate physical transmission processes, this pattern is consistent with forms of coordination that are less spatially targeted and potentially operate over longer timescales [39]. Such modes of interaction may facilitate persistent neural states required for higher-level cognitive functions, including reasoning and emotional regulation [42]. Notably, this hierarchical transition aligns with regional molecular and synaptic landscapes [30]. Cortical areas exhibiting stronger Syn_TU_ dominance show higher expression of synapse-related genes—particularly those involved in chemical transmission and receptor activity—alongside enriched receptor diversity. These features may provide a flexible molecular substrate that supports functional coordination less fully constrained by tractography [35]. Together, the observed shift from predominantly wiring-aligned synergistic interactions toward more flexible, modulatory forms of synchrony highlights an organizational principle that enables the cortex to balance physiological stability with the complexity and adaptability required for advanced human cognition.

Consistent with prior work on functional fingerprinting [7], the preferential concentration of Syn_TU_ in higher-order cognitive regions—where structural–functional decoupling is most pronounced [19]—suggests that this tract-underexplained dimension captures aspects of functional organization relevant to individual specificity. By partitioning the functional connectome, our framework isolates Syn_TE_ as a conserved, structurally constrained backbone while revealing individual-specific functional patterns within Syn_TU_ that are otherwise masked by shared anatomical architecture. From a broader perspective, this implies that individual-specific functional organization is preferentially expressed along dimensions that are weakly constrained by anatomical wiring, rather than within the structurally conserved backbone of the connectome.

Although this framework provides a new perspective on the organization of functional connectivity, several limitations warrant consideration. First, while the model explicitly separates tract-mediated and non-tract-mediated components, residual overlap between Syn_TE_ and Syn_TU_ may persist, potentially limiting the precision with which their unique contributions can be quantified. Future methodological advances aimed at more tightly constraining the model could further improve this separation. Second, the present analyses are restricted to resting-state data. Extending this framework to task-based paradigms will be important for understanding how the balance between structurally constrained and flexible interactions is dynamically reconfigured during cognition. Third, although Syn_TU_ is associated with multiscale biological features, the model does not explicitly incorporate underlying physiological mechanisms. Linking model parameters to in vivo measures such as receptor density or synaptic processes will be an important step toward a more biologically grounded and generative framework. Finally, the current study is cross-sectional. Longitudinal investigations across development and aging, as well as applications in clinical contexts, will be necessary to determine whether the enhanced sensitivity of Syn_TU_ to individual variability can be leveraged for tracking disease progression or treatment response.

In summary, our framework disentangles functional connectivity into two complementary components: Syn_TE_, which reflects rigid anatomical constraints supporting stable global integration, and Syn_TU_, which captures flexible, non-tract modulation driving region-specific adaptation. This dual-layered perspective advances our understanding of large-scale brain network architecture by reconciling structurally constrained and flexible modes of brain organization. By highlighting the unique sensitivity of Syn_TU_ to individual variability, this work provides a powerful new tool for decoding complex cognition, and offers both a generalizable framework and a new conceptual perspective for understanding how diverse biological factors jointly shape functional connectivity, with support from cross-species validation.

## 4 Methods

### 4.1 Multimodal Datasets

See Supplementary Methods for detailed descriptions of multimodal data acquisition and preprocessing (including MRI, PET, MEG, gene expression, and *in vitro* autoradiography). Briefly, the diverse datasets utilized in this study were curated from multiple independent resources: (1) structural and functional MRI data were obtained from the Human Connectome Project (HCP) [43], the Boston Adolescent Neuroimaging of Depression and Anxiety (BANDA) cohort [44], and a cross-species marmoset dataset from the Marmoset Brain Mapping (MBM) Project [32]; (2) group-level PET maps, accessed via the neuromaps toolbox [38], were integrated to capture neurotransmitter receptors and transporters [38], aerobic glycolysis (glycolytic index) [45], and synaptic density [46]; (3) MEG data used for deriving cortical temporal dynamics were sourced from the HCP S900 release [43]; (4) regional microarray gene expression data were acquired from the Allen Human Brain Atlas (AHBA) [47]; and (5) *in vitro* quantitative autoradiography data characterizing neurotransmitter receptor densities were derived from post-mortem human brains [48].

### 4.2 Connectomes

For the human datasets, the whole brain was parcellated into 100 regions using the Schaefer-100 atlas [29]. For each region, the mean time series was obtained by averaging the BOLD signal across all voxels within the parcel. To enhance the stability of functional connectivity (FC) estimation, multiple scanning sessions per subject were concatenated into a single extended time series. Static FC matrices were then computed as Pearson correlation coefficients between all pairs of regional time series [5].

For the marmoset dataset, cortical parcellation was performed using the MBMv4 atlas [32]. Regional mean time series were extracted in the same manner as for the human data, and FC matrices were computed using Pearson correlation. Structural connectivity was obtained from the group-level data provided by the dataset.

For dynamic functional connectivity (dFC), a phase based approach was adopted following previous studies [49]. At each time point *t*, the dFC between region *n* and region *m* was calculated according to the following equation:

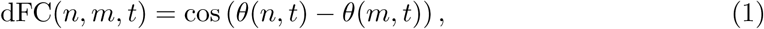

where *θ*(*n, t*) denotes the instantaneous phase of region *n* at time *t*, which was obtained by applying the Hilbert transform to the bandpass filtered time series (0.04–0.08 Hz). From the dFC, functional connectivity dynamics (FCD) and probabilistic metastable substates (PMS) were further derived [49, 50]. FCD was computed by vectorizing the upper triangular portion of the dFC matrix at each time point, and then calculating the cosine similarity between these vectors across all time points, resulting in a *T × T* FCD matrix that reflects the temporal dynamics of functional connectivity patterns. PMS were estimated as follows: first, the leading eigenvector of the dFC matrix was extracted at each time point to capture the dominant connectivity mode. These eigenvectors were then clustered using k-means clustering to identify recurrent connectivity states. Each time point was assigned to one of the clusters, and the probability of occurrence of each state—defined as the proportion of time points assigned to that cluster—was computed to obtain the probabilistic metastable substates.

For structural connectivity (SC), probabilistic tractography was performed for each subject using fiber orientation distribution (iFOD2) algorithm implemented in MRtrix3 [51, 52]. The Schaefer-100 atlas was used to define 100 cortical parcels, and SC matrices were generated by counting the number of streamlines between each pair of regions. Each individual’s SC matrix was normalized to a maximum value of 1. To ensure positive values while achieving numerical scaling, a power transformation with exponent 0.1 was applied (rather than logarithmic transformation) [14, 53]. The diagonal elements were then set to zero.

### 4.3 Linearized parametric large-scale brain model

To simulate macroscale neural dynamics, our large-scale brain model was built upon a previously established dynamic mean-field (DMF) model introduced by Deco et al [12]. Within this whole-brain network, the macroscopic neuronal activity of the *i*-th cortical region was described by a nonlinear differential equation:

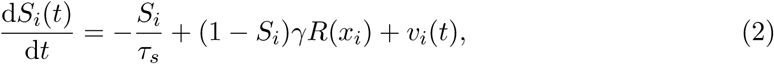

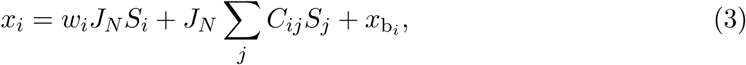

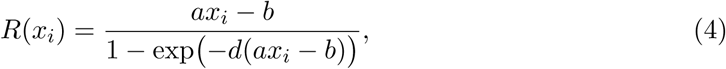

where *S_i_* denotes the synaptic gating variable, *x_i_* represents the total input to brain region *i*, *R*(*x_i_*) is the mean firing rate of region *i*, *C_ij_* indicates the large-scale inter-regional coupling between regions *i* and *j* that is constrained by structural connectivity, *w_i_* is the self-feedback coefficient of region *i*, and 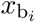 represents the baseline non-cortical input. The dynamical parameters were set as follows [12]: *τ_s_* = 0.1 s*, γ* = 0.641*, a* = 270 n/C*, b* = 108 Hz*, d* = 0.154 s*, J_N_* = 0.3.

Following previous large-scale modeling studies, *w_i_* and 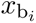 were treated as heterogeneous parameters across regions [14, 26]. Recent work has emphasized the importance of both co-operative and competitive interactions between brain regions at the macroscale. Although long-range projections have traditionally been viewed as primarily excitatory, emerging evidence has also supported the existence of inhibitory pathways in large-scale connectivity [54, 55]. To account for these complex regional interactions across the brain, functional connections that showed consistently negative correlations across four resting-state scans were identified and incorporated into the large-scale coupling matrix *C_ij_* as stable mutual inhibition. Experimental evidence has demonstrated that the low-frequency components of the local field potential (LFP) are strongly correlated with BOLD signal fluctuations [56, 57]. In addition, theoretical work by Demirtas et al. indicated that the analytical solution of the hemodynamic model approximates a low-pass filter [25]. Therefore, the model variable *x*, which represents the macroscopic LFP input, was bandpass filtered between 0.01–0.1 Hz to simulate large-scale BOLD signals for the computation of both static and dynamic functional connectivity.

As in prior work by kong et al. [26], *w_i_* and 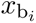 were expressed as linear combinations of local myelin content (T1w/T2w ratio) and the principal functional connectivity gradient to capture macroscale hierarchies:

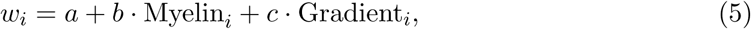

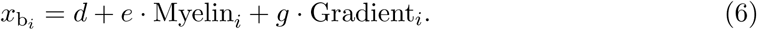

For the large-scale inter-regional coupling *C_ij_*, we embedded competitive interactions derived from empirical negative functional connections into the structural connectome. We then introduced three global scaling factors to modulate whole-brain dynamics: *G*_1_ for within-network connections, *G*_2_ for between-network connections, and *G*_3_ specifically for negative connections.

As computing correlations between nodes of this high-dimensional large-scale brain model driven by stochastic noise, as defined in Eqs. (2)-(4), is computationally expensive and sensitive to random fluctuations, we analytically linearized the system around its fixed point. We first perform a first-order Taylor expansion of the activation function *R*(*x*) around the steady-state input 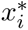 and synaptic gating variable 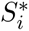, resulting in the following expression:

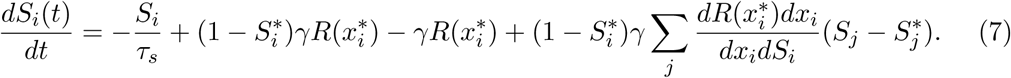

After simplification, the system can be rewritten as:

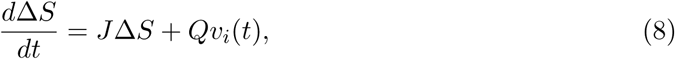

where *J* is the Jacobian matrix and *Q* is the perturbation coupling matrix. Specifically:

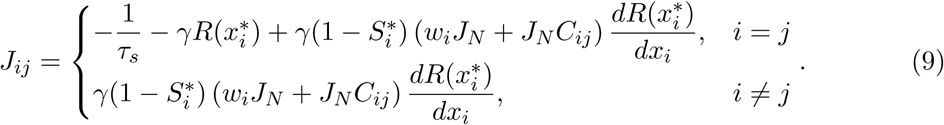

The cross-spectral density of the system in the frequency range 0.01–0.1 Hz was computed analytically via the transfer matrix method. The covariance matrix of the input variable *x* is given by:

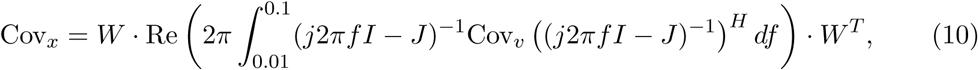

where Cov*_v_* represents the covariance matrix of the background stochastic noise *v* across brain regions, and the element *W_ij_* in the projection matrix *W* is defined as:

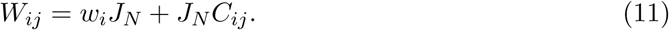

Finally, the covariance matrix was normalized to produce the correlation coefficient matrix, which captures the inter-node correlations of the system and corresponds to the simulated static functional connectivity:

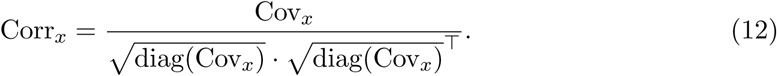

### 4.4 Model Parameter Optimization

Based on previous theoretical work, functional correlations among brain regions are thought to be largely shaped by structural connectivity [12, 26]. We therefore optimized model heterogeneity parameters such that, given structural connectivity, the optimized large-scale model could maximally explain empirical functional connectivity. Specifically, in Eq. (10), Cov*_v_* was set as a diagonal matrix, assuming that brain regions were independent in the absence of structural connectivity. Under this assumption, the linearized parametric large-scale brain model for each participant contained nine free parameters. In the absence of biological priors, the model involves 203 free parameters. To verify the computational feasibility of our optimization framework in a high-dimensional parameter space, we performed a validation by fitting all 203 parameters for the first 50 participants in the HCP dataset (Fig. S1A, B). This confirmed that the optimization process remains robust even without the constraints of biological priors. However, the parsimonious nine-parameter version was ultimately selected for the main analyses to prevent overfitting and ensure the interpretability and robustness of the model across the entire cohort.

These parameters were optimized individually using the covariance matrix adaptation evolution strategy (CMA-ES) [58], with the loss function defined as the mean absolute error (MAE) between the simulated functional connectivity generated by the model and the empirical functional connectivity. Each optimization was repeated ten times, and the solution with the smallest loss was selected as the final optimized parameter set. This approach improved convergence and reduced the risk of trapping in local minima.

### 4.5 Solving for the Covariance Matrix

Even after parameter optimization with *C_v_* constrained to be diagonal, the simulated functional connectivity did not fully capture the empirical functional connectivity, leaving residual error. Inspired by Pang et al. [20], we assumed that large-scale empirical functional connectivity may not be entirely determined by structural connectivity, and therefore allowed *C_v_* to take a non-diagonal form. This accounts for the possibility that, even in the absence of structural connectivity, brain regions may exhibit intrinsic synchrony. Accordingly, we directly solved for the *C_v_* that best reproduced the empirical functional connectivity using the optimized model as the generative framework. We solved for *C_v_* based on Eq. (10). Notably, changes in cross-spectral density within the frequency band 0.01–0.1 Hz were negligible relative to their absolute values and exhibited monotonic behavior. Therefore, we approximated the spectral density using its value at 0.055 Hz:

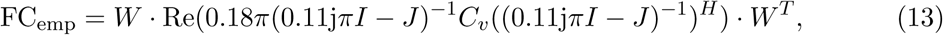

where *J* is the Jacobian matrix and *W* is the projection matrix.

Further inspection of the equation above showed that the real component of the system response dominated the imaginary component by approximately four orders of magnitude. Hence, the imaginary contribution was neglected, leading to the simplified formulation:

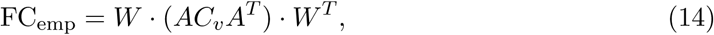

where *A* denotes the real component of the system transfer matrix. This formulation enabled us to estimate the covariance matrix of the background stochastic noise in the optimized model, thereby accounting for the residual error after parameter optimization.

### 4.6 Extraction of Tract-Explainable and Underexplained Synchrony

Based on the optimized model with the solved *C_v_*, we further disentangled the respective contributions of structural connectivity and background noise covariance. To isolate the component explained by structural connectivity, *C_v_* was constrained to a diagonal matrix, yielding simulated functional connectivity denoted as Syn_TE_. Conversely, to characterize the component not explained by structural connectivity, structural connectivity was set to zero while retaining the solved non-diagonal *C_v_*, yielding simulated functional connectivity denoted as Syn_TU_.

### 4.7 Graph-Based Network Measures

We treated Syn_TE_ and Syn_TU_ as two continuous functional synchrony networks and compared them in terms of their graph-theoretical properties. Our analysis focused on quantifying two fundamental principles of brain network organization: network segregation and global integration.

To quantify network segregation, we computed the weighted modularity coefficient, which evaluates the goodness with which a network is partitioned into distinct functional sub-groups [10, 27]. For fully connected, weighted functional networks, we employed the generalized asymmetric modularity metric *Q*^∗^:

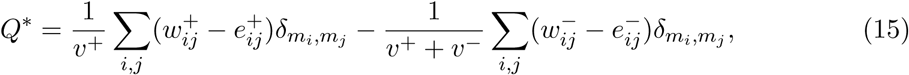

where 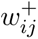 and 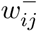 denote the positive and negative connection weights, *v*^+^ and *v*^−^ are the respective sums of all positive or negative weights in the network, 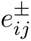 represents the chance-expected within-module weights, and the Kronecker delta *δ_m__i,mj_* equals 1 if nodes *i* and *j* belong to the same module, and 0 otherwise. This asymmetric formulation is preferred as it accounts for the intrinsically unequal biological roles of positive and negative correlations in resting-state functional networks.

To quantify global integration, we measured the average connection diversity. Unlike the traditional participation coefficient (which is based on the Simpson index and suffers from a variable upper bound), we utilized the normalized Shannon entropy, as it provides a consistent upper bound of 1 across different network partitions [27]. Because resting-state functional networks contain both positive and negative correlations with inherently unequal biological roles, we computed the asymmetric connection diversity 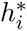 for each node *i*. First, the connection diversity for positive and negative weights 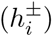 is defined as:

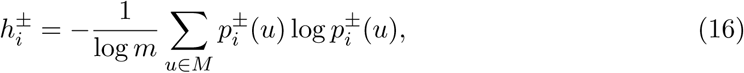

where *m* is the total number of modules, and 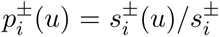 represents the proportion of node *i*’s total positive or negative strength 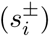 that connects to nodes within module *u*. To systematically integrate both components while preserving the primary role of positive connections, the final regional connection diversity 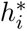 is computed by scaling the negative diversity contribution:

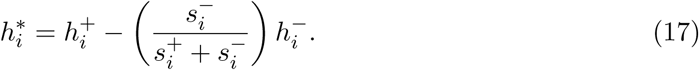

To yield a global macroscale metric, the regional diversity scores 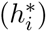 were subsequently averaged across all nodes to obtain the mean connection diversity for both the Syn_TE_ and Syn_TU_ components.

### 4.8 Functional Gradient

We first computed the group-averaged FC, Syn_TE_, and Syn_TU_, setting their diagonal elements to zero to remove self-connections. Diffusion map embedding, implemented in the BrainSpace toolbox [59], was then applied to each matrix to derive gradient components.

### 4.9 Explained–Underexplained Synchrony Consistency

We extracted the column vectors of Syn_TU_-Syn_TE_ for each brain region. For a given region, the two vectors respectively represented its functional synchrony pattern with all other regions in the two distinct modes. Inspired by the concept of structure–function coupling [15], we then computed the Spearman correlation coefficient between the corresponding column vectors of Syn_TU_ and Syn_TE_ for each region. We defined this coefficient as the Explained–Underexplained Synchrony Consistency, whose magnitude reflects the degree of concordance between the two functional synchrony modes of that region.

### 4.10 Gradient of Syn_TU_-Syn_TE_ Relative Importance

Relative importance was quantified using the Underexplained–Minus–Explained Rank, defined as the difference between the nodal strength rank in Syn_TU_ and that in Syn_TE_. This metric follows the framework proposed by Luppi et al. for the gradient of redundancy-to-synergy relative importance [28]. Specifically, we first computed nodal strength for both Syn_TU_ and Syn_TE_, defined as the sum of the absolute values of all connections incident to each ROI. Regions were then ranked separately according to their nodal strength in Syn_TU_ and Syn_TE_, with higher-strength regions assigned higher ranks. For each region, the Syn_TE_ rank was subtracted from the Syn_TU_ rank, yielding the UMER gradient. This gradient ranges from negative values—indicating regions more strongly ranked in Syn_TE_—to positive values, indicating regions that are relatively more prominent in Syn_TU_.

The same procedure was also repeated for network edges rather than nodes. Edge weights were used to rank connections separately according to Syn_TE_ and Syn_TU_, and the difference between rankings was then calculated. This resulted in a single connectivity matrix representing the relative importance of edges.

### 4.11 Meta-analysis

We conducted a meta-analysis, following approaches used in previous studies by Margulies et al. and Preti et al. [3, 19], to characterize the behavioral and cognitive relevance of the regional relationship between Syn_TE_ and Syn_TU_. Using the NeuroSynth framework, we performed a topic-based association analysis to link spatial variations in this relationship to a predefined set of psychological and behavioral terms. To this end, we derived regional relationship scores that summarize how Syn_TU_ and Syn_TE_ are related within each cortical region. These relationship scores were then divided into five percentile bins, yielding twenty binary masks that partitioned the cortical surface according to different regimes of the Syn_TU_–Syn_TE_ relationship. These masks were subsequently used as inputs for the meta-analysis, following the same procedure described above. Finally, we assessed the association between each mask and 24 selected psychological and behavioral terms using NeuroSynth, allowing us to identify cognitive and behavioral domains systematically associated with regional variations in the relationship between Syn_TU_ and Syn_TE_.

### 4.12 Gene Enrichment Analysis

We first computed the Pearson correlation between the regional expression level of each gene and the regional Syn_TE_–Syn_TU_ interaction gradient scores. To control for multiple comparisons across the transcriptome, the resulting *P* -values were corrected using the Benjamini–Hochberg false discovery rate (FDR) procedure. Genes exhibiting significant positive or negative associations (FDR *<* 0.05) were then segregated into two distinct sets for subsequent analysis. Comprehensive gene set enrichment analysis was performed on these filtered gene sets using GSEApy via the Enrichr module [60]. To capture a broad biological context, we simultaneously interrogated multiple databases, encompassing three Gene Ontology (GO) domains—Biological Processes (BP), Molecular Functions (MF), and Cellular Components (CC) [61]—as well as the Reactome database for signaling and metabolic pathways [62].

### 4.13 Common and Individual Functional Connectivity

To disentangle subject-specific network features from shared population-level signals, we applied a spatial filtering approach based on principal component analysis (PCA), following the connectome caricatures framework [31]. Specifically, an eigendecomposition was performed on the group-concatenated functional time series to extract the top six dominant spatial eigenvectors, which represent large-amplitude coactivation patterns shared across the cohort. By projecting the empirical time series of each subject onto this group-level manifold, we partitioned the data into two distinct dimensions: a “common” functional connectivity matrix reconstructed from these first six principal components, reflecting canonical network architecture, and an “individual” functional connectivity matrix derived from the remaining residual components. This latter projection effectively filters out shared global variance, thereby emphasizing idiosyncratic, subject-specific functional configurations that are typically masked by large-amplitude coactivations.

### 4.14 Partial Least Squares (PLS) Analysis

Partial least squares (PLS) analysis was used to investigate the multivariate relationships between functional synchrony components (Syn_TE_, Syn_TU_, and empirical FC) and complex cognitive abilities [63]. PLS is a robust, data-driven multivariate statistical technique designed to identify mutually orthogonal linear combinations of two distinct blocks of variables that maximally covary with each other. In the present analysis, prior to applying PLS, the multidimensional behavioral data underwent strict quality control and preprocessing. To enforce normality and mitigate the influence of outliers, each of the ten cognitive task scores was first transformed using a rank-based inverse normal transformation (RINT) [64]. Subsequently, the effects of age and sex were regressed out from the normalized cognitive scores using ordinary least squares (OLS) regression. To identify the latent variables, the resulting residual behavioral matrix, denoted as *Y* (subjects *×* cognitive tasks), and the vectorized upper-triangular functional connectivity matrix, denoted as *X* (subjects *×* functional edges), were both normalized column-wise (i.e., z-scored), and a singular value decomposition (SVD) was applied to the cross-covariance matrix *R* = *X^T^ Y* as follows:

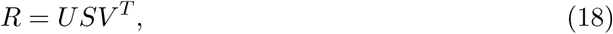

where *U* and *V* are orthonormal matrices containing the left and right singular vectors (often termed spatial and cognitive weights, respectively), and *S* is the diagonal matrix of singular values. Each corresponding column of the *U* and *V* matrices defines a latent variable (LV). The singular values (*s_k_*) in *S* indicate the amount of cross-covariance explained by each LV. We specifically utilized the absolute covariance, defined as 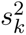, to evaluate the effect size of each LV. Subject-specific brain and cognitive scores for each LV, which demonstrate the extent to which each subject expresses the weighted patterns identified by the latent variables, were computed by projecting the original standardized data onto their PLS-derived weights:

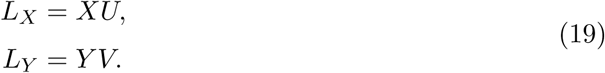

The overall statistical significance of each latent variable (LV) was assessed using a non-parametric permutation test [65]. Specifically, the rows of the behavioral matrix *Y* were randomly shuffled to disrupt subject identity, and the SVD-PLS was re-computed 1,000 times. This iterative process generated an empirical null distribution of the absolute covariance 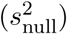 for each LV. The *p*-value was then calculated as the proportion of permuted 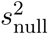 values that were greater than or equal to the actual 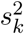, yielding a robust and highly conservative estimate of significance.

### 4.15 Statistical Analysis

The statistical significance of within-individual differences between Syn_TE_ and Syn_TU_—such as variations in participation coefficient, modularity, and associations with multiscale cortical features—was determined using paired-samples two-tailed *t* -tests. Effect sizes for these comparisons were estimated using Cohen’s measure of the standardized mean difference, *d*.

To rigorously control for the confounding effects of spatial autocorrelation in brain maps, we assessed the statistical significance of cortical topography correlations using a spatial permutation framework (‘spin test’) [66]. This approach generates null distributions by applying 5,000 random spherical rotations to the cortical surface, thereby preserving the intrinsic spatial covariance structure of the data. This non-parametric test was explicitly applied to evaluate the significance of associations between the Syn_TE_–Syn_TU_ interaction gradient and spatial maps such as the sensorimotor-association axis, cortical developmental expansion, and intrinsic neural timescales.

Furthermore, to verify that the spatial coupling between EUSC and UMER reflects true biological organization rather than mathematical redundancy, we conducted a group-level permutation test. This involved generating 1,000 surrogate networks by randomly shuffling empirical connectivity matrices while preserving the degree distribution. Finally, where multiple hypotheses were tested simultaneously—specifically for gene and pathway enrichment analyses and evaluations across multiple MEG frequency bands—*P* values were corrected for multiple comparisons using the false discovery rate (FDR) based on the Benjamini–Hochberg method [67].

## Data availability

The raw and minimally preprocessed data used in this study are available from publicly accessible repositories. Resting-state fMRI, structural MRI, diffusion MRI, and behavioral data from both the Human Connectome Project (HCP) young adult cohort and the Boston Adolescent Neuroimaging of Depression and Anxiety (BANDA) dataset are available via the Connectome Coordination Facility (https://www.humanconnectome.org/). All marmoset-related data, including the resting-state fMRI and structural connectivity matrices, are publicly available through the Marmoset Brain Mapping Project at www.marmosetbrainmapping.org/data.html. Microarray gene expression data were obtained from the Allen Human Brain Atlas (AHBA; http://human.brain-map.org/).

Neurotransmitter receptor and transporter PET maps, group-averaged synaptic density ([11C]UCB-J PET) maps, glycolytic index (GI) maps, as well as MEG-derived intrinsic timescales and source-level power estimates from the HCP S900 release, are all available via the neuromaps toolbox (https://netneurolab.github.io/neuromaps/). Microstructure profile covariance (MPC) matrices were obtained from the Microstructure-Informed Connectomics (MICA-MICs) database (https://portal.conp.ca/dataset?id=projects/mica-mics).

## Code availability

The preprocessing codes for fiber tracking and structural connectivity are available at https://github.com/sina-mansour/UKB-connectomics. Upon acceptance of this manuscript, the specific codes for our model will be made publicly available on GitHub at https://github.com/GuoLab-UESTC. The code used for NeuroSynth meta-analysis is freely available at https://www.github.com/gpreti/GSP_StructuralDecouplingIndex. Spatial autocorrelation-preserving null models were implemented using the codebase available at https://github.com/netneurolab/shafiei_megdynamics. Gene set enrichment analyses were performed using the GSEApy package (https://gseapy.readthedocs.io/).

## Acknowledgments

We sincerely thank Prof. Songting Li, Prof. Yunliang Zang, and Dr. Lin Jiang for their insightful discussions and valuable suggestions on this work. This work was supported in part by the Brain Science and Brain-like Intelligence Technology-National Science and Technology Major Project (Grant No. 2022ZD0208500), the National Key Research and Development Program of China (Grant No. 2023YFF1204200), the Lingang Laboratory (Grant No. LGL-1987), the National Natural Science Foundation of China (Grant No. 62572485), and the Sichuan Science and Technology Program (Grant Nos. 2024NSFJQ0004, DQ202410, and 2024NSFTD0032). Data were provided in part by the Human Connectome Project, WU-Minn Consortium (Principal Investigators: David Van Essen and Kamil Ugurbil; 1U54MH091657) funded by the 16 NIH Institutes and Centers that support the NIH Blueprint for Neuroscience Research; and by the McDonnell Center for Systems Neuroscience at Washington University.

## Author Contributions

**J.L.** and **D.G.** conceived the research and designed the study. **J.L.** and **X.Z.** were responsible for data preprocessing and software development. **Y.X.** and **Y.X. (Xu)** assisted with data curation and statistical analysis. **C.Z.**, **Y.W.**, **D.G.** and **D.Y.** provided methodology guidance and critical resources. **D.Y.**, **D.G.** and **Y.W.** acquired the fundings. **J.L.** wrote the original draft. **D.G.**, **D.Y.** and **C.Z.** supervised the project and revised the manuscript. All authors reviewed and approved the final manuscript.

## Competing Interests

The authors declare no competing interests.

## Notes

### Competing Interest Statement

The authors have declared no competing interest.

### Summary of Updates

Both the maintext and supplemental files updated

## References

[1] Buckner, R. L., Krienen, F. M. & Yeo, B. T. T. Opportunities and limitations of intrinsic functional connectivity MRI. Nature Neuroscience 16, 832–837 (2013).

[2] Brookes, M. J. et al. Measuring functional connectivity using MEG: Methodology and comparison with fcMRI. NeuroImage 56, 1082–1104 (2011).

[3] Margulies, D. S. et al. Situating the default-mode network along a principal gradient of macroscale cortical organization. Proceedings of the National Academy of Sciences of the United States of America 113, 12574–12579 (2016).

[4] Fox, M. D. & Raichle, M. E. Spontaneous fluctuations in brain activity observed with functional magnetic resonance imaging. Nature Reviews Neuroscience 8, 700–711 (2007).

[5] Biswal, B., Yetkin, F., Haughton, V. & Hyde, J. Functional Connectivity in the Motor Cortex of Resting Human Brain Using Echo-Planar Mri. Magnetic Resonance in Medicine 34, 537–541 (1995).

[6] Shen, X. et al. Using connectome-based predictive modeling to predict individual behavior from brain connectivity. Nature Protocols 12, 506–518 (2017).

[7] Finn, E. S. et al. Functional connectome fingerprinting: Identifying individuals using patterns of brain connectivity. Nature Neuroscience 18, 1664–1671 (2015).

[8] Dubois, J., Galdi, P., Paul, L. K. & Adolphs, R. A distributed brain network predicts general intelligence from resting-state human neuroimaging data. Philosophical Transactions of the Royal Society B: Biological Sciences 373 (2018).

[9] Yuste, R. From the neuron doctrine to neural networks. Nature Reviews Neuroscience 16, 487–497 (2015).

[10] Rubinov, M. & Sporns, O. Complex network measures of brain connectivity: Uses and interpretations. NeuroImage 52, 1059–1069 (2010).

[11] Zhang, F. et al. Quantitative mapping of the brain’s structural connectivity using diffusion MRI tractography: A review. NeuroImage 249, 118870 (2022).

[12] Deco, G. et al. Resting-State Functional Connectivity Emerges from Structurally and Dynamically Shaped Slow Linear Fluctuations. Journal of Neuroscience 33, 11239–11252 (2013).

[13] Honey, C. J. et al. Predicting human resting-state functional connectivity from structural connectivity. Proceedings of the National Academy of Sciences of the United States of America 106, 2035–2040 (2009).

[14] Zhang, S. et al. In vivo whole-cortex marker of excitation-inhibition ratio indexes cortical maturation and cognitive ability in youth. Proceedings of the National Academy of Sciences of the United States of America 121, e2318641121.

[15] Fotiadis, P. et al. Structure–function coupling in macroscale human brain networks. Nature Reviews Neuroscience 25, 688–704 (2024).

[16] Gu, Z., Jamison, K. W., Sabuncu, M. R. & Kuceyeski, A. Heritability and interindividual variability of regional structure-function coupling. Nature Communications 12, 4894 (2021).

[17] Zamani Esfahlani, F., Faskowitz, J., Slack, J., Mišić, B. & Betzel, R. F. Local structure-function relationships in human brain networks across the lifespan. Nature Communications 13, 2053 (2022).

[18] Baum, G. L. et al. Development of structure–function coupling in human brain networks during youth. Proceedings of the National Academy of Sciences of the United States of America 117, 771–778 (2020).

[19] Preti, M. G. & Van De Ville, D. Decoupling of brain function from structure reveals regional behavioral specialization in humans. Nature Communications 10, 4747 (2019).

[20] Pang, J. C. et al. Geometric constraints on human brain function. Nature 618, 566–574 (2023).

[21] Richiardi, J. et al. Correlated gene expression supports synchronous activity in brain networks. Science 348, 1241–1244 (2015).

[22] Markello, R. D. et al. Neuromaps: Structural and functional interpretation of brain maps. Nature Methods 19, 1472–1479 (2022).

[23] Feng, G. et al. Spatial and temporal pattern of structure–function coupling of human brain connectome with development. eLife 13, RP93325 (2024).

[24] Deco, G. et al. Dynamical consequences of regional heterogeneity in the brain’s transcriptional landscape. Science Advances 7, eabf4752 (2021).

[25] Demirtaş, M., et al. Hierarchical Heterogeneity across Human Cortex Shapes Large-Scale Neural Dynamics. Neuron 101, 1181–1194.e13 (2019).

[26] Kong, X. et al. Sensory-motor cortices shape functional connectivity dynamics in the human brain. Nature Communications 12, 6373 (2021).

[27] Rubinov, M. & Sporns, O. Weight-conserving characterization of complex functional brain networks. NeuroImage 56, 2068–2079 (2011).

[28] Luppi, A. I. et al. A synergistic core for human brain evolution and cognition. Nature Neuroscience 25, 771–782 (2022).

[29] Schaefer, A. et al. Local-Global Parcellation of the Human Cerebral Cortex from Intrinsic Functional Connectivity MRI. Cerebral Cortex 28, 3095–3114 (2018).

[30] Goulas, A. et al. The natural axis of transmitter receptor distribution in the human cerebral cortex. Proceedings of the National Academy of Sciences of the United States of America 118, e2020574118 (2021).

[31] Rodriguez, R. X., Noble, S., Camp, C. C. & Scheinost, D. Connectome caricatures remove large-amplitude coactivation patterns in resting-state fmri to emphasize individual differences. Nature Neuroscience 1–11 (2025).

[32] Tian, X. et al. An integrated resource for functional and structural connectivity of the marmoset brain. Nature Communications 13, 7416 (2022).

[33] Agnati, L. F., Guidolin, D., Guescini, M., Genedani, S. & Fuxe, K. Understanding wiring and volume transmission. Brain Research Reviews 64, 137–159 (2010).

[34] Shine, J. M., Aburn, M. J., Breakspear, M. & Poldrack, R. A. The modulation of neural gain facilitates a transition between functional segregation and integration in the brain. eLife 7, e31130 (2018).

[35] Shine, J. M. Neuromodulatory Influences on Integration and Segregation in the Brain. Trends in Cognitive Sciences 23, 572–583 (2019).

[36] Marder, E. Neuromodulation of neuronal circuits: back to the future. Neuron 76, 1–11 (2012).

[37] Veening, J. G. & Barendregt, H. P. The regulation of brain states by neuroactive substances distributed via the cerebrospinal fluid; a review. Cerebrospinal Fluid Research 7, 1 (2010).

[38] Hansen, J. Y. et al. Mapping neurotransmitter systems to the structural and functional organization of the human neocortex. Nature Neuroscience 25, 1569–1581 (2022).

[39] Murray, J. D. et al. A hierarchy of intrinsic timescales across primate cortex. Nature Neuroscience 17, 1661–1663 (2014).

[40] Owen, L. L. W. & Manning, J. R. High-level cognition is supported by information-rich but compressible brain activity patterns. Proceedings of the National Academy of Sciences of the United States of America 121, e2400082121.

[41] Buckner, R. L. & Krienen, F. M. The evolution of distributed association networks in the human brain. Trends in Cognitive Sciences 17, 648–665 (2013).

[42] Honey, C. J. et al. Slow cortical dynamics and the accumulation of information over long timescales. Neuron 76, 423–434 (2012).

[43] Van Essen, D. C. et al. The Human Connectome Project: A data acquisition perspective. NeuroImage 62, 2222–2231 (2012).

[44] Hubbard, N. et al. Brain function and clinical characterization in the boston adolescent neuroimaging of depression and anxiety study. NeuroImage: Clinical 27, 102240 (2020).

[45] Vaishnavi, S. N. et al. Regional aerobic glycolysis in the human brain. Proceedings of the National Academy of Sciences of the United States of America 107, 17757–17762 (2010).

[46] Finnema, S. J. et al. Kinetic evaluation and test-retest reproducibility of [11C]UCB-J, a novel radioligand for positron emission tomography imaging of synaptic vesicle glycoprotein 2A in humans. Journal of Cerebral Blood Flow and Metabolism: Official Journal of the International Society of Cerebral Blood Flow and Metabolism 38, 2041–2052 (2018).

[47] Hawrylycz, M. J. et al. An anatomically comprehensive atlas of the adult human brain transcriptome. Nature 489, 391–399 (2012).

[48] Zilles, K. & Palomero-Gallagher, N. Multiple transmitter receptors in regions and layers of the human cerebral cortex. Frontiers in Neuroanatomy 11, 78 (2017).

[49] Hansen, E. C. A., Battaglia, D., Spiegler, A., Deco, G. & Jirsa, V. K. Functional connectivity dynamics: Modeling the switching behavior of the resting state. NeuroImage 105, 525–535 (2015).

[50] Cabral, J. et al. Cognitive performance in healthy older adults relates to spontaneous switching between states of functional connectivity during rest. Scientific Reports 7, 5135 (2017).

[51] Tournier, J.-D. et al. Mrtrix3: A fast, flexible and open software framework for medical image processing and visualisation. NeuroImage 202, 116137 (2019).

[52] Tournier, J. D., Calamante, F., Connelly, A., et al. Improved probabilistic streamlines tractography by 2nd order integration over fibre orientation distributions, Vol. 1670 (Stockholm, 2010).

[53] Park, B.-y., et al. Differences in subcortico-cortical interactions identified from connectome and microcircuit models in autism. Nature Communications 12, 2225 (2021).

[54] Luppi, A. I. et al. Competitive interactions shape mammalian brain network dynamics and computation. Nature Neuroscience 1–19 (2026).

[55] Melzer, S. et al. Long-Range–Projecting GABAergic Neurons Modulate Inhibition in Hippocampus and Entorhinal Cortex. Science 335, 1506–1510 (2012).

[56] Li, J. M., Acland, B. T., Brenner, A. S., Bentley, W. J. & Snyder, L. H. Relationships between correlated spikes, oxygen and LFP in the resting-state primate. NeuroImage 247, 118728 (2022).

[57] Pan, W.-J., Thompson, G. J., Magnuson, M. E., Jaeger, D. & Keilholz, S. Infraslow LFP correlates to resting-state fMRI BOLD signals. NeuroImage 74, 288–297 (2013).

[58] Lozano, J. A., Larrañaga, P., Inza, I., Bengoetxea, E. & Kacprzyk, J. (eds) Towards a New Evolutionary Computation Vol. 192 of Studies in Fuzziness and Soft Computing (Springer, Berlin, Heidelberg, 2006).

[59] Vos de Wael, R., et al. BrainSpace: A toolbox for the analysis of macroscale gradients in neuroimaging and connectomics datasets. Communications Biology 3, 103 (2020).

[60] Fang, Z., Liu, X. & Peltz, G. GSEApy: A comprehensive package for performing gene set enrichment analysis in Python. Bioinformatics 39, btac757 (2023).

[61] Carbon, S. et al. The Gene Ontology Resource: 20 years and still GOing strong. Nucleic Acids Research 47, D330–D338 (2019).

[62] Fabregat, A. et al. The Reactome Pathway Knowledgebase. Nucleic Acids Research 46, D649–D655 (2018).

[63] McIntosh, A. R. & Lobaugh, N. J. Partial least squares analysis of neuroimaging data: applications and advances. NeuroImage 23, S250–S263 (2004).

[64] Puth, M.-T., Neuhäuser, M. & Ruxton, G. D. Effective use of Pearson’s product–moment correlation coefficient. Animal Behaviour 93, 183–189 (2014).

[65] Nichols, T. E. & Holmes, A. P. Nonparametric permutation tests for functional neuroimaging: A primer with examples. Human Brain Mapping 15, 1–25 (2002).

[66] Alexander-Bloch, A. F. et al. On testing for spatial correspondence between maps of human brain structure and function. NeuroImage 178, 540–551 (2018).

[67] Benjamini, Y. & Hochberg, Y. Controlling the False Discovery Rate: A Practical and Powerful Approach to Multiple Testing. Journal of the Royal Statistical Society: Series B (Methodological) 57, 289–300 (1995).

