## Supplementary material for "Tract-explainable and underexplained synchrony play complementary roles in the functional organization of the brain": SI

### Supplementary Information

#### 1 Supplementary Methods

##### 1.1 Neuroimaging Datasets

Two independent human neuroimaging cohorts were included in this study: the Human Connectome Project (HCP) [1] and the Boston Adolescent Neuroimaging of Depression and Anxiety (BANDA) [2].

From the HCP cohort, 1,012 subjects were selected who had completed four minimally preprocessed resting-state fMRI (rs-fMRI) sessions, along with corresponding structural MRI (sMRI), diffusion tensor imaging (DTI), and behavioral data.

From the BANDA cohort, 202 subjects were included, comprising 63 healthy controls, 62 individuals with depression, and 77 with anxiety. All selected subjects had four minimally preprocessed rs-fMRI sessions and matched sMRI, DTI, and behavioral data.

In addition to the human cohorts, we included a non-human primate dataset consisting of 24 marmosets with both diffusion MRI (dMRI) and resting-state fMRI (rs-fMRI) data [3]. This cross-species dataset enabled us to assess the reproducibility and generalizability of the proposed framework across species.

Details of the imaging protocols and data acquisition parameters for each dataset are available in the original publications and on the respective data-sharing platforms.

##### 1.2 Data processing

For both human cohorts, minimally preprocessed rs-fMRI data provided by the respective repositories were used. The HCP and BANDA datasets were processed using the minimal preprocessing pipelines [4], which included correction for spatial artifacts and distortions, surface reconstruction, cross-modal registration, and alignment to standard space, followed by denoising with ICA-FIX [5]. Global signal regression was not included in the preprocessing pipeline for either dataset, and an additional 0.01–0.1 Hz bandpass filter was applied to retain neural-related low-frequency fluctuations.

Diffusion MRI data from both human cohorts underwent identical preprocessing steps: $b_0$ intensity normalization, correction for echo-planar imaging (EPI) distortions and eddy current-induced distortions, motion correction, gradient nonlinearity correction, and affine registration to the T1-weighted (T1w) anatomical space [6].

For the marmoset dataset, preprocessed rs-fMRI data (regrBMWC0) were obtained from a publicly available resource. This preprocessing pipeline included regression of linear and quadratic trends, band-pass filtering (0.01–0.1 Hz), and nuisance regression of motion parameters (including derivatives), white matter signals, cerebrospinal fluid signals, and motion-censor regressors.

Volumes with excessive motion were censored based on framewise displacement thresholds, and data were subsequently normalized to the MBMv3 template space.

##### 1.3 Gene Expression Data

We obtained regional microarray expression data from six post-mortem human brains (1 female, ages 24–57 years, mean $\pm$ SD: $42.5 \pm 13.4$) provided by the Allen Human Brain Atlas (AHBA) [7]. We processed the data using the abagen toolbox with the Schaefer-100 volumetric atlas in MNI space [8].

First, we reannotated microarray probes using updated probe information and discarded probes without a valid Entrez ID. We further filtered probes based on their expression intensity relative to background noise, removing those with intensity below background in 50% of samples across donors. For genes indexed by multiple probes, we selected the probe with the highest differential stability, defined as the average Spearman correlation of expression modes across all pairs of donors. We updated the MNI coordinates of tissue samples using nonlinear registration with Advanced Normalization Tools (ANTs; `https://github.com/chrisfilo/alleninf`). To enhance spatial coverage, we mirrored tissue samples bilaterally across hemispheres. We assigned samples to atlas parcels if their MNI coordinates were within 2 mm of a parcel. For regions not directly assigned a sample, we estimated expression values using distance-weighted interpolation from the nearest tissue samples of each donor and averaged values across voxels. We discarded tissue samples that could not be mapped to any parcel.

To address inter-subject variation, we normalized expression values across genes within each sample using a robust sigmoid function, where each value was rescaled relative to the sample median and interquartile range, and subsequently mapped to the unit interval. The same normalization procedure was applied across samples. Finally, we averaged samples assigned to the same region within donors and then across donors, yielding a parcellated regional expression matrix of 15,631 genes. Gene expression similarity (GS) was defined as the Pearson correlation coefficient between regional gene expression vectors. The depression-related gene co-expression network was calculated using disease priors related to Major Depressive Disorder provided by the ENIGMA tools [9].

##### 1.4 PET Data

Neurotransmitter receptor and transporter data were obtained from group-level volumetric positron emission tomography (PET) maps provided by Hansen et al. [10], comprising 19 neurotransmitter receptors and transporters across nine neurotransmitter systems: serotonin (5-$HT_{1a}$, 5-$HT_{1b}$, 5-$HT_{2a}$, 5-$HT_4$, 5-$HT_6$, 5-HTT), histamine ($H_3$), dopamine ($D_1$, $D_2$, DAT), norepinephrine (NET), acetylcholine ($\alpha_4\beta_2$, $M_1$, VAChT), cannabinoid ($CB_1$), opioid (MOR), glutamate ($mGluR_5$, NMDA), and GABA ($GABA_A$/bz). For each receptor map, we extracted the mean receptor density within regions defined by the Schaefer-100 parcellation, followed by z-score normalization across regions. Receptor similarity (RS) was then defined as the Pearson correlation coefficient between regional receptor density vectors across brain regions.

##### 1.5 Glycolytic Index Data

We obtained group-averaged maps of the glycolytic index (GI) from Vaishnavi et al. [11], based on positron emission tomography (PET) scans of 33 healthy adults. GI reflects regional aerobic glycolysis, calculated as the deviation of empirically measured glucose consumption from that predicted by oxygen metabolism. Specifically, we derived GI by regressing the regional cerebral metabolic rate of glucose on the cerebral metabolic rate of oxygen, measured using three O-PET tracers (water, carbon monoxide, oxygen). We then extracted regional GI values using the Schaefer-100 parcellation.

##### 1.6 Synapse Density Data

We used group-level maps of synaptic density provided by Finnema et al. [12], derived from positron emission tomography (PET) with the radioligand [$^{11}$C]UCB-J. This tracer binds selectively to synaptic vesicle glycoprotein 2A (SV2A), thereby providing an *in vivo* measure of synaptic density. The dataset was based on scans acquired in 76 healthy adults (45 males, mean ± SD: 48.9 ± 18.4 years) on an HRRT PET camera over 90 minutes post-injection. Non-displaceable binding potential ($BP_{ND}$) was estimated using the SRTM2 model with the centrum semiovale as reference tissue. We parcellated the group-level synapse density maps into 100 cortical regions using the Schaefer-100 atlas.

##### 1.7 Receptor Entropy

We quantified receptor entropy based on neurotransmitter receptor densities obtained from *in vitro* quantitative autoradiography. Data included 15 receptor types across multiple neurotransmitter systems: glutamate (AMPA, NMDA, kainate), GABA ($GABA_A$, $GABA_A$/BZ, $GABA_B$), acetylcholine (muscarinic $M_1$, $M_2$, $M_3$, nicotinic $\alpha_4\beta_2$), noradrenaline ($\alpha_1$, $\alpha_2$), serotonin ($5\text{-}HT_{1A}$, $5\text{-}HT_2$), and dopamine ($D_1$). Receptor densities were measured across 44 cytoarchitectonically defined cortical areas in three postmortem brains (age range: 72–77 years, two males) [13]. We retained 43 areas with complete receptor profiles and estimated receptor diversity following the method of Goulas et al. [14]. Specifically, for each area, we constructed a normalized receptor profile vector $P$, where each entry denotes the density of a given receptor divided by its maximum density across all areas. Receptor entropy was then computed as the Shannon entropy $H$ of $P$, providing a measure of receptor diversity.

We mapped receptor entropy values onto the Schaefer-100 parcellation using the atlas transformation procedure provided by Hansen et al. [10]. When multiple cytoarchitectonic areas corresponded to the same Schaefer-100 parcel, we averaged their entropy values.

##### 1.8 Microstructure Profile Covariance

The microstructure profile covariance (MPC) matrix was obtained from a group-level template provided by the Microstructure-Informed Connectomics (MICA-MICs) database [15], which captures inter-regional similarities in cortical microstructural profiles.

##### 1.9 Cortical Developmental Expansion

We obtained the map of relative cortical surface area expansion between term birth and adulthood from the study by Hill et al. [16]. Briefly, the expansion map was derived by examining the fractional surface area at each cortical location. For each hemisphere of every individual, a map of fractional surface area was calculated by normalizing the surface area of each fiducial surface tile by the total cortical surface area, excluding the non-cortical medial wall. The map of relative areal expansion was then constructed by calculating the ratio between the group-averaged fractional area map of adults and that of term infants. To align with the functional parcellation used in the present study, the vertex-wise expansion map was parcellated into 100 cortical regions using the Schaefer-100 atlas.

##### 1.10 Time-series Features

We derived brain dynamics features from magnetoencephalography (MEG) data obtained from the Human Connectome Project (HCP S900 release, $n = 33$) [1]. The preprocessing pipeline followed the procedures described in the Brainstorm online tutorials and in previous work by Shafiei et al [17, 18]. For each subject, we reconstructed MEG time series at the source level using the Schaefer-100 parcellation [19]. We then computed a total of 6,880 time-series features for each time point using the hctsa toolbox and assigned them to their corresponding brain regions [20]. After feature computation, we z-score normalized the feature matrices at the individual level and constructed a group-average region-by-feature matrix by averaging the normalized subject-level matrices. We applied PCA to this matrix and retained the first principal component—referred to as the Timescore—as a principal gradient of cortical temporal dynamics [17].

We also obtained intrinsic timescales and source-level power estimates across six canonical frequency bands—delta (2–4 Hz), theta (5–7 Hz), alpha (8–12 Hz), beta (15–29 Hz), low gamma (30–59 Hz), and high gamma (60–90 Hz)—from publicly available data via Neuromaps [21]. Intrinsic timescales were estimated using the FOOOF algorithm [22], which decomposes the power spectra into periodic (oscillatory) and aperiodic (1/f-like) components by fitting the power spectral density. Specifically, the algorithm identifies oscillatory peaks (periodic component), the "knee parameter" $k$ that controls the bend in the aperiodic component, and the aperiodic exponent $\chi$. We then used the knee parameter to calculate the "knee frequency" as $f_k = k^{1/\chi}$, which corresponds to the frequency where a bend occurs in the power spectrum. Finally, we estimated the intrinsic timescale $\tau$ as:

$$\tau = \frac{1}{2\pi f_k}. \tag{1}$$

##### 1.11 Geometry-Constrained Functional Connectivity

To evaluate the specific contribution of cortical geometry to functional synchrony, we generated a simulated functional connectivity matrix driven purely by the brain's spatial structure. This approach is grounded in neural field theory (NFT), which posits that macroscopic neural dynamics are fundamentally shaped by the physical contours of the cortex, propagating as waves across the continuous cortical surface [23].

The spatiotemporal dynamics of this wave propagation are governed by an isotropic damped wave equation. Projected onto an orthogonal spatial basis formed by the geometric eigenmodes of the cortex, the equation is given by:

$$\sum_{j=1}^{N}\left[\frac{1}{\gamma_s^2}\frac{\partial^2}{\partial t^2}+\frac{2}{\gamma_s}\frac{\partial}{\partial t}+1+r_s^2\lambda_j\right]\phi_j(t)\psi_j(\mathbf{r})=\sum_{j=1}^{N}q_j(t)\psi_j(\mathbf{r}). \tag{2}$$

In this formulation, $\psi_j(\mathbf{r})$ represents the $j$-th spatial eigenmode derived directly from the cortical geometry, and $\lambda_j$ is its corresponding eigenvalue. For the wave dynamics, $\gamma_s$ denotes the damping rate, $r_s$ is the spatial length scale of wave propagation, $q_j(t)$ represents the time-varying amplitude obtained from the mode decomposition of empirical data, $\phi_j(t)$ is the temporal component of mode $j$, and $N$ is the total number of modes.

By directly adopting the optimal parameters and geometric eigenmodes provided by Pang et al. [23], we solved this differential equation for the temporal components $\phi_j(t)$. We then simulated the spatiotemporal neural activity across the cortex by combining these temporal components with their corresponding spatial eigenmodes $\psi_j(\mathbf{r})$. Finally, a simulated functional connectivity matrix was derived by computing the Pearson correlation of these simulated regional time series. We regarded this simulated matrix as the geometry-constrained functional connectivity, representing a baseline mode of functional coordination that arises exclusively from the brain's physical geometry.

##### 1.12 Gaussian Linear Diffusion Model

To examine the generality of our proposed framework, we also utilized a Gaussian linear diffusion (GLD) model to simulate the dynamics of local node in the large-scale brain models [24]. Assuming a local excitation-inhibition balance where regional inputs interact with the current activation level, the collective dynamics of the $i$-th neural population follows a Gaussian linear process and can be described as follows:

$$\dot{x}_i=-a_i x_i+\sum_{j=1}^{N}b_i C_{ij}(x_j-x_i)+\sqrt{2}\xi_i, \tag{3}$$

where $x_i$ represents the neural activity (or macroscopic firing activity) of the $i$-th brain region. The parameters $a_i$ and $b_i$ denote regional heterogeneity, representing the local relaxation rate and the coupling sensitivity of region $i$, respectively. Specifically, $a_i$ governs the intrinsic decay of neural activity in each region, while $b_i$ scales the influence of incoming inputs from the rest of the network. $C_{ij}$ indicates the inter-regional coupling constrained by structural connectivity, and $\xi_i$ is Gaussian white noise with zero mean and unit variance.

#### 2 Supplementary Figures

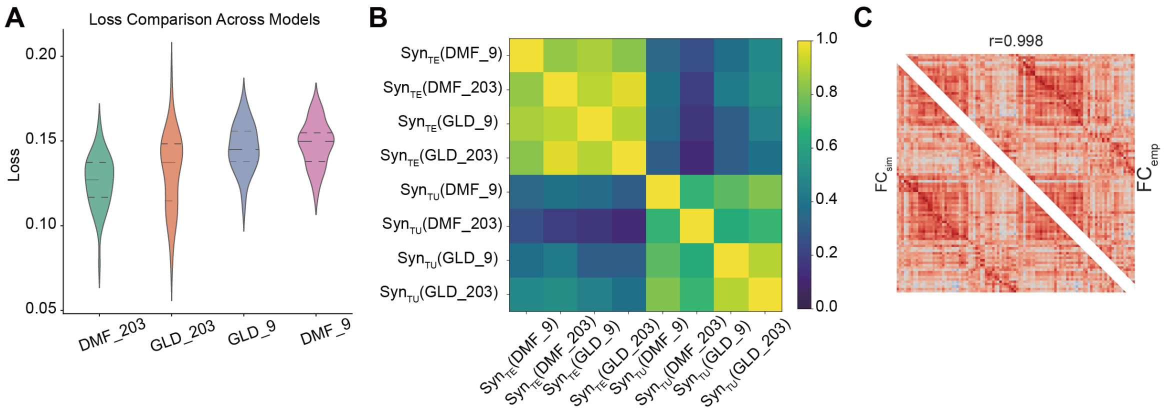


**Fig. S1 Evaluation of the extraction framework.** **(A)** Comparison of the final loss, quantified by the Mean Absolute Error (MAE), during the first-stage individual-level optimization. The evaluation contrasts the dynamic mean-field (DMF) model and the Gaussian linear diffusion (GLD) model across two parameterization scales: a parsimonious model with 9 free parameters (constrained by biological priors) and an unconstrained model with 203 free parameters (allowing for full regional heterogeneity). Data are shown for the first 50 participants of the HCP dataset. **(B)** Assessment of the similarity between $\mathrm{Syn}_{\mathrm{TE}}$ and $\mathrm{Syn}_{\mathrm{TU}}$ estimates obtained across various models and parameter settings, demonstrating the robustness of the decomposition. **(C)** Reconstruction performance for a randomly selected participant, quantified by the similarity between the empirical functional connectivity ($\mathrm{FC}_{\mathrm{emp}}$) and the simulated functional connectivity ($\mathrm{FC}_{\mathrm{sim}}$) generated from the extracted synchrony components.

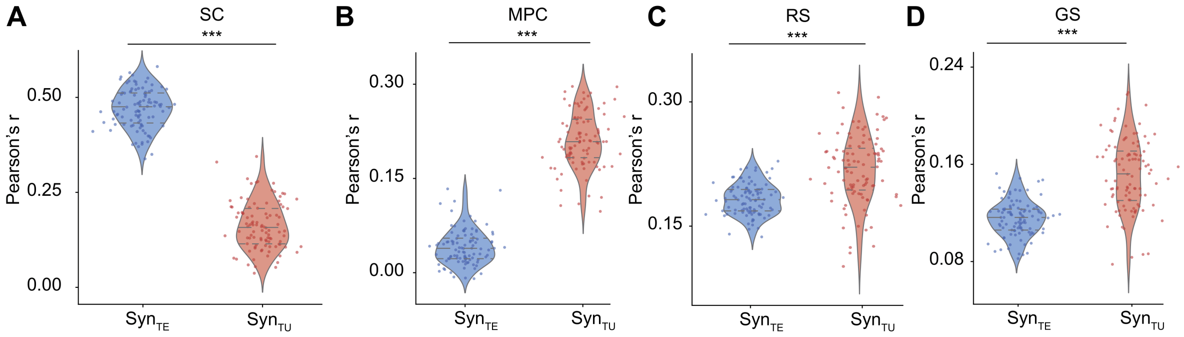


**Fig. S2 Differential relationships of Syn$_{\mathrm{TE}}$ and Syn$_{\mathrm{TU}}$ with tractography-derived structural connectivity and multiscale cortical similarity features without distance regression.** **(A)** Comparison of the association between synchrony components and structural connectivity (SC), where the correlation for Syn$_{\mathrm{TE}}$ remained significantly stronger than that for Syn$_{\mathrm{TU}}$ (paired-samples $t$-test, $t(99) = 34.631$, $d = 3.463$, $***p < 0.001$). **(B–D)** Assessment of the associations with non-tractography cortical features, showing that the correlations between Syn$_{\mathrm{TE}}$ and **(B)** microstructure profile covariance (MPC) ($t(99) = -28.809$, $d = -2.881$, $***p < 0.001$), **(C)** receptor similarity (RS) ($t(99) = -7.263$, $d = -0.726$, $***p < 0.001$), and **(D)** gene similarity (GS) ($t(99) = -10.269$, $d = -1.027$, $***p < 0.001$) were all significantly weaker than those observed for Syn$_{\mathrm{TU}}$.

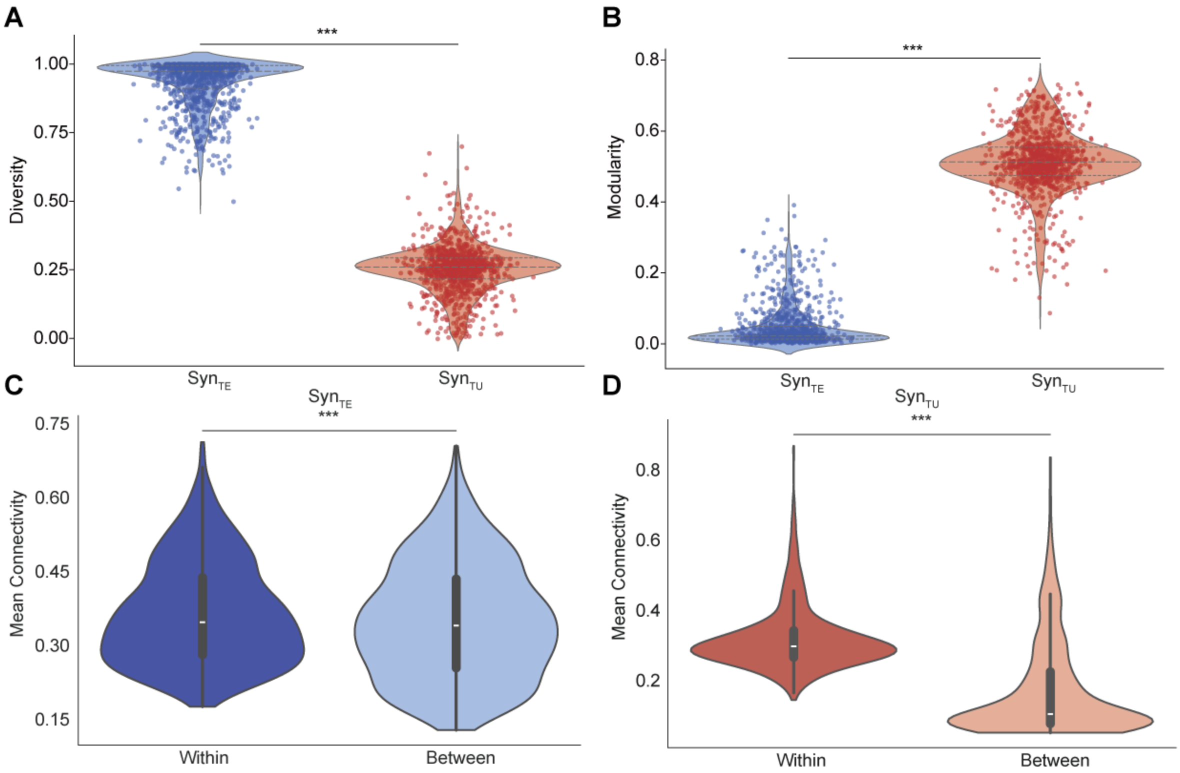


**Fig. S3 Different functional synchrony patterns exhibit distinct network properties at the individual level. (A)** The average connection diversity of $\mathrm{Syn}_{\mathrm{TE}}$ was significantly higher than that of $\mathrm{Syn}_{\mathrm{TU}}$ ($t(1011) = 197.516$, $d = 6.209$, $***p < 0.001$). **(B)** The modularity of $\mathrm{Syn}_{\mathrm{TE}}$ was significantly lower than that of $\mathrm{Syn}_{\mathrm{TU}}$ ($t(1011) = -172.942$, $d = -5.436$, $***p < 0.001$). **(C)** $\mathrm{Syn}_{\mathrm{TE}}$ was significantly higher within resting-state subnetworks (RSNs) than between them ($t(1011) = 21.599$, $d = 0.679$, $***p < 0.001$). **(D)** $\mathrm{Syn}_{\mathrm{TU}}$ was significantly higher within RSNs than between them ($t(1011) = 59.086$, $d = 1.857$, $***p < 0.001$).

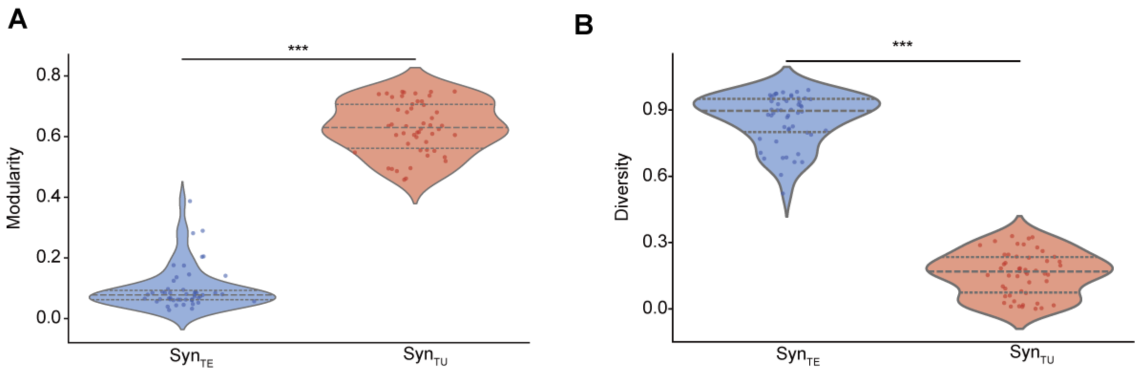


**Fig. S4 Robustness of topological properties across higher-resolution parcellations.** Results were validated using a finer parcellation scheme (Schaefer-400 atlas) in a random subsample of 50 participants. **(A)** The modularity of $\mathrm{Syn}_{\mathrm{TU}}$ was significantly higher than that of $\mathrm{Syn}_{\mathrm{TE}}$ (paired-samples $t$-test, $t(49) = 32.348$, $d = -4.575$, $^{***}p < 0.001$). **(B)** The average connection diversity of $\mathrm{Syn}_{\mathrm{TE}}$ was significantly higher than that of $\mathrm{Syn}_{\mathrm{TU}}$ (paired-samples $t$-test, $t(49) = 32.733$, $d = 4.629$, $^{***}p < 0.001$).

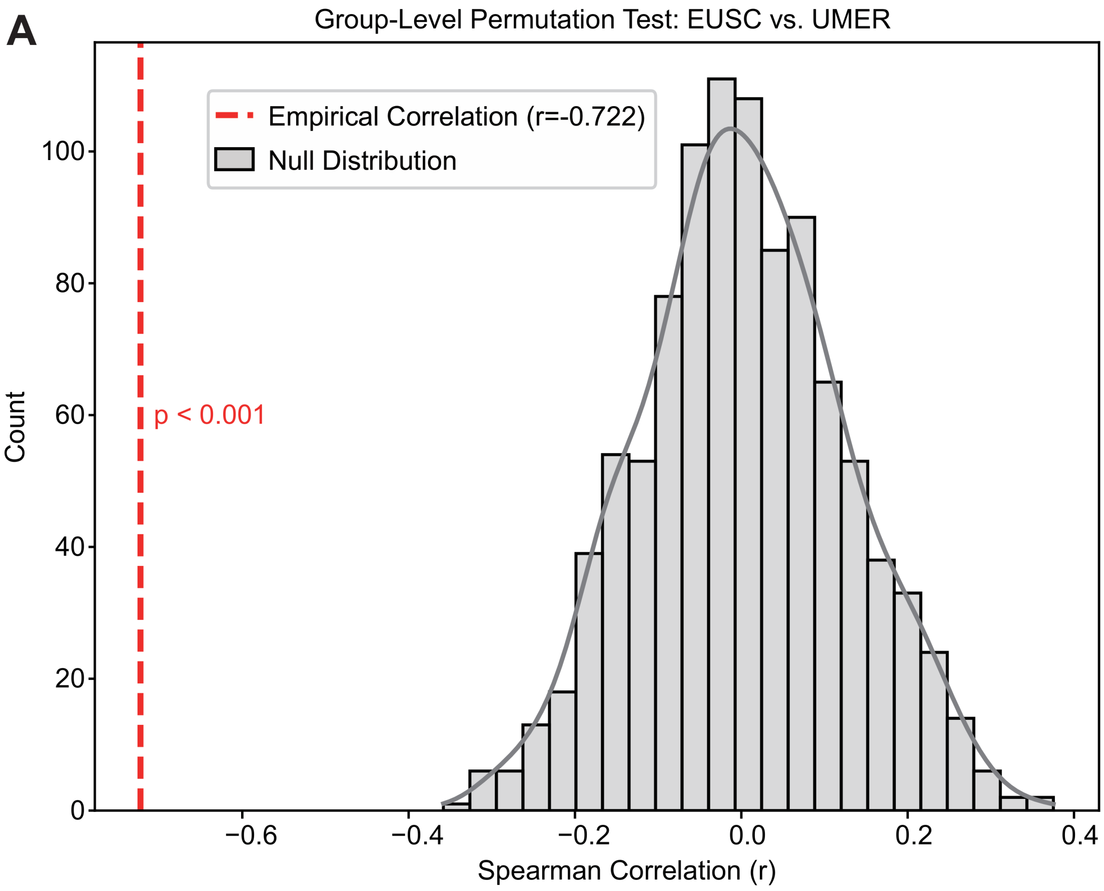


**Fig. S5 Group-level permutation test for the coupling between EUSC and UMER.** **(A)** To assess the statistical significance of the spatial correlation between the Observed Explained–Underexplained Synchrony Consistency (EUSC) and the Underexplained–Minus–Explained Rank (UMER), a group-level permutation test was performed ($n = 1,000$ permutations). For each permutation, the nodal indices of the $\mathrm{Syn_{TU}}$ were independently shuffled for each participant before recomputing the group-averaged metrics. The red dashed line indicates the empirical Spearman correlation ($r = -0.722$), which sits far outside the null distribution (gray histogram) generated from the permuted data ($p < 0.001$). This result demonstrates that the tight coupling between $\mathrm{Syn_{TE}}$–$\mathrm{Syn_{TU}}$ spatial dissociation and their functional dominance shift is not a mathematical artifact of the nodal properties but represents a non-trivial organizational feature of the brain.

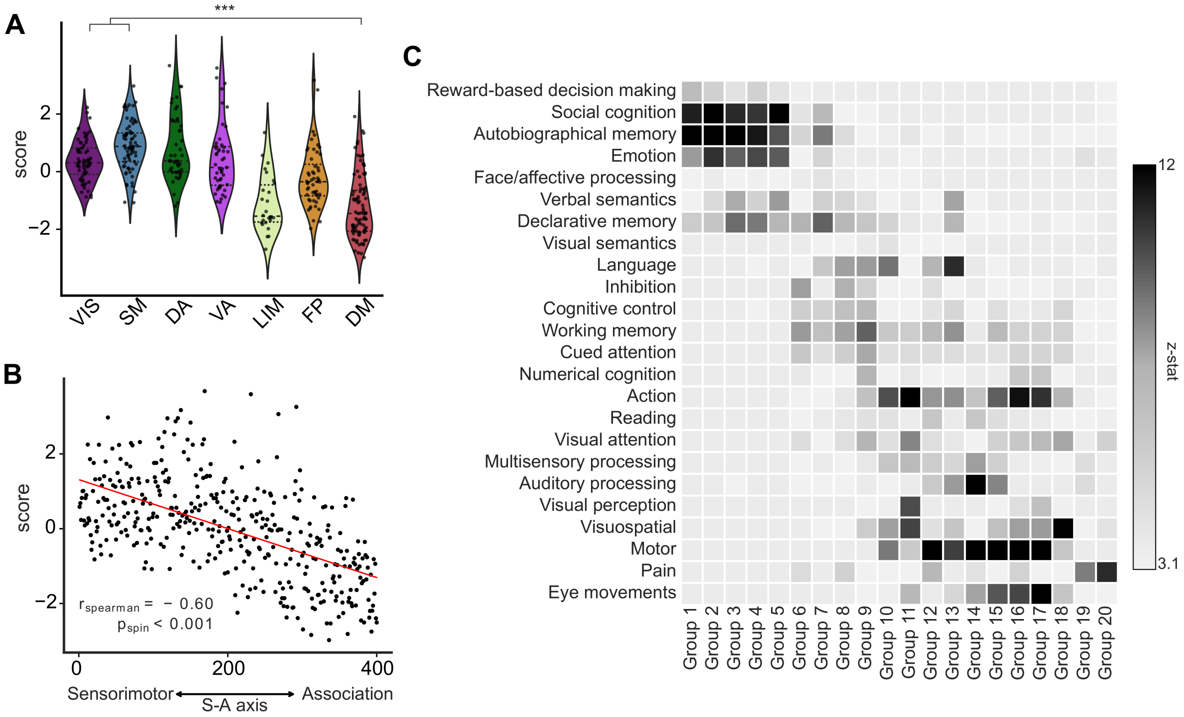


**Fig. S6 The $\text{Syn}_{\text{TE}}$–$\text{Syn}_{\text{TU}}$ interaction gradient reflects the functional hierarchy of the brain across the Schaefer 400 parcellation. (A)** The distribution of PC1 scores across the seven resting-state networks is shown. Each dot represents a brain region; the central thick dashed line indicates the median, and the thin dashed lines denote the first and third quartiles. The default mode (DM) network exhibits significantly lower PC1 scores than the visual (VIS) and somatomotor (SM) networks (***$p < 0.001$). **(B)** The association between PC1 and regional position along the sensorimotor–association (S–A) cortical axis is quantified using Spearman correlation ($r_{\text{spearman}} = -0.60$, $p_{\text{spin}} < 0.001$, two-tailed). **(C)** NeuroSynth-based meta-analysis of PC1. Brain regions are ranked by PC1 and binned into 20 equally sized groups along the $x$-axis, while the $y$-axis lists cognitive and behavioral terms from NeuroSynth. Heatmap intensity reflects the mean task-activation $z$-scores within each bin (analysis pipeline adapted from Margulies et al.).

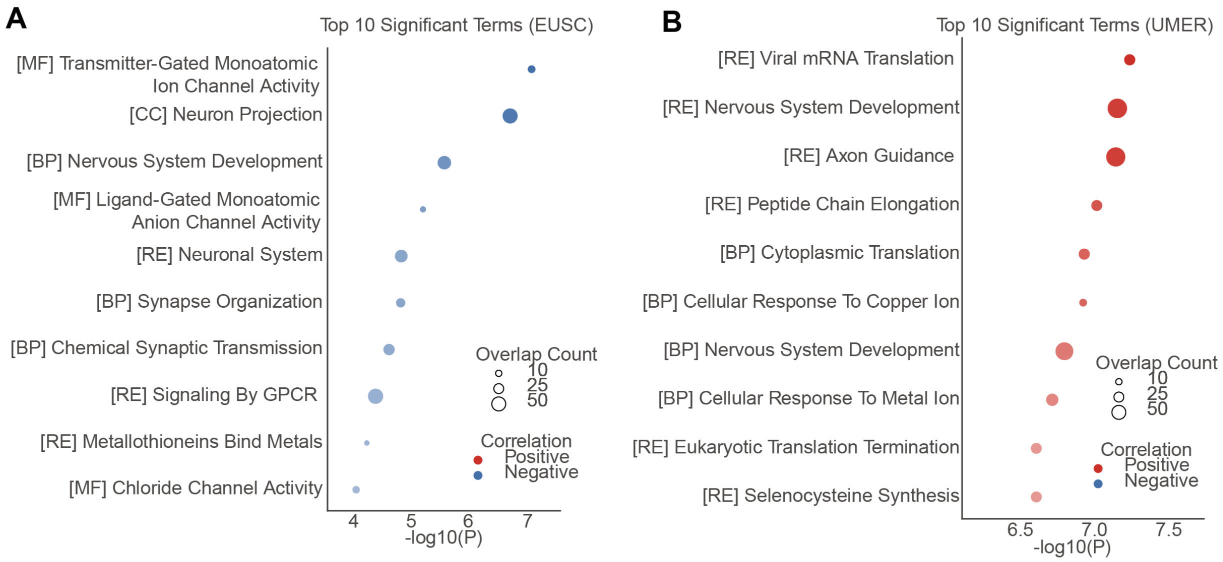


**Fig. S7 Comprehensive gene set enrichment analyses associated with the individual spatial metrics EUSC and UMER.** **(A)** Comprehensive gene set enrichment analysis associated with the Explained–Underexplained Synchrony Consistency (EUSC). To capture a broader biological context, the analysis integrates multiple databases: Gene Ontology (GO), Biological Processes ([BP]), Molecular Functions ([MF]), Cellular Components ([CC]), and Reactome pathways ([RE]). The top ten significantly enriched terms across all combined datasets are shown. Bubble color indicates the direction of correlation (red = positive; blue = negative), and bubble size reflects the number of overlapping genes contributing to each term. **(B)** Comprehensive gene set enrichment analysis associated with the Underexplained–Minus–Explained Rank (UMER). The analysis integrates the same multiple databases ([BP], [MF], [CC], and [RE]) to display the top ten significantly enriched terms. Bubble color and size follow the same conventions as in **(A)**. Statistical significance for all analyses was assessed using FDR correction via the Benjamini–Hochberg method.

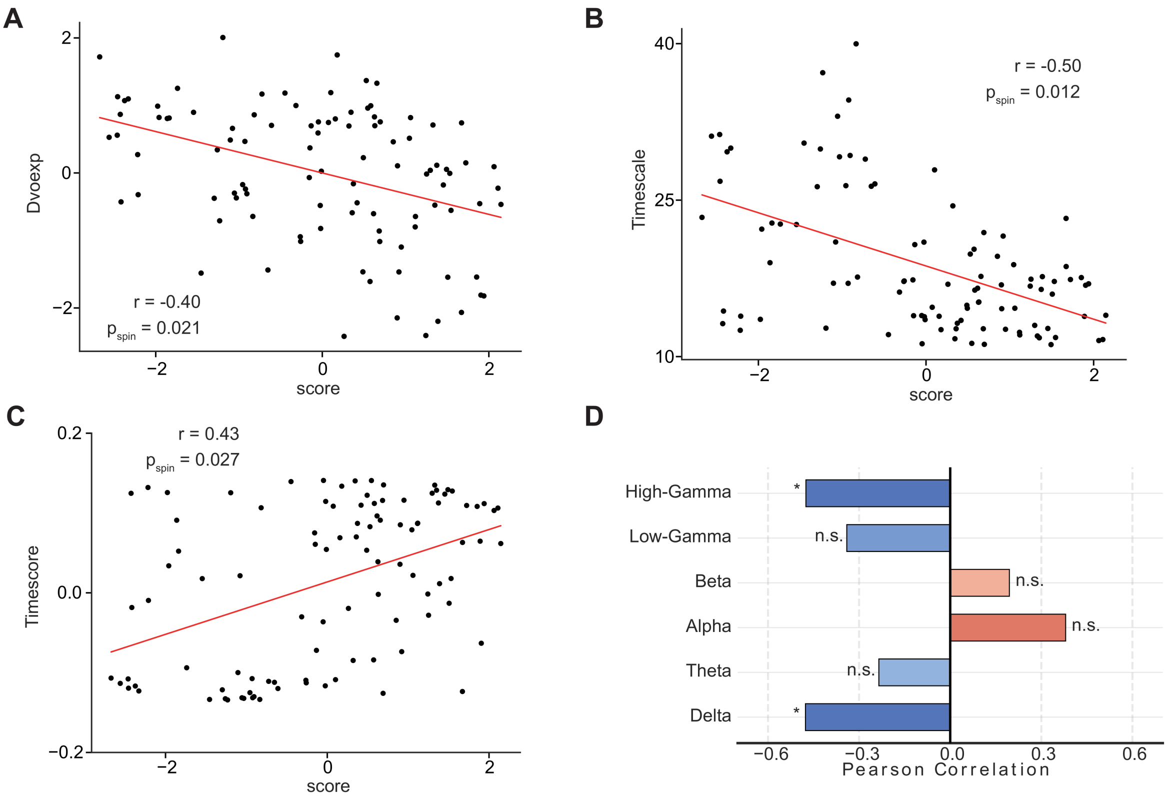


**Fig. S8 Association of the $\text{Syn}_{\text{TE}}$–$\text{Syn}_{\text{TU}}$ interaction gradient with various forms of cortical regional heterogeneity. (A)** The association between gradient scores and cortical developmental expansion was quantified using Pearson correlation ($r = -0.40$, $p_{\text{spin}} = 0.021$, two-tailed). **(B)** The association between gradient scores and intrinsic neural timescales was quantified using Pearson correlation ($r = -0.50$, $p_{\text{spin}} = 0.012$, two-tailed). **(C)** The association between gradient scores and timescore was quantified using Pearson correlation ($r = -0.50$, $p_{\text{spin}} = 0.027$, two-tailed). **(D)** The association between gradient scores and the mean power across various frequency bands was assessed using two-tailed spin tests with FDR correction.

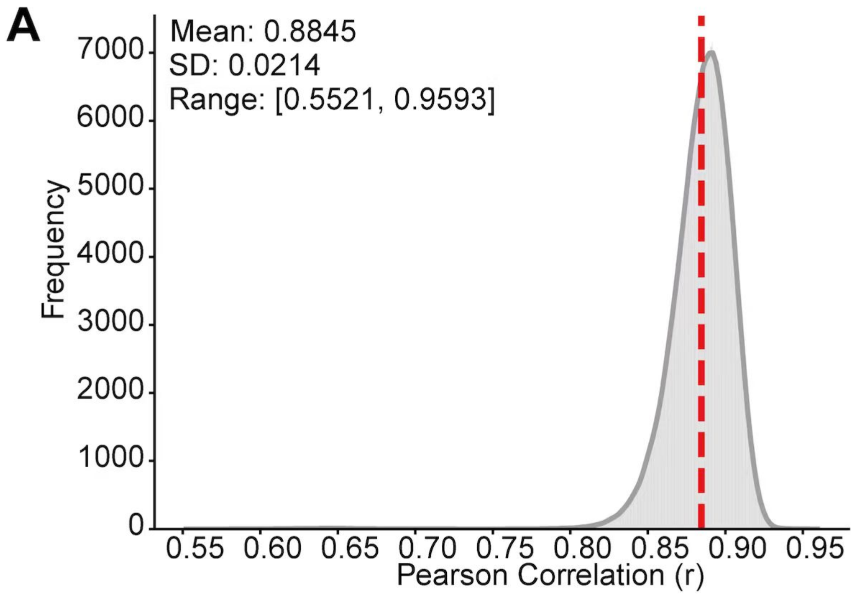


**Fig. S9 Inter-individual consistency of empirical structural connectivity. (A)** Distribution of pairwise inter-subject similarity (Pearson correlation) for empirical structural connectivity profiles across the study cohort. The empirical SC exhibits exceptionally high reproducibility across individuals, with a mean inter-individual correlation of $r = 0.8845$ (SD = 0.0214, range: [0.5521, 0.9593]).

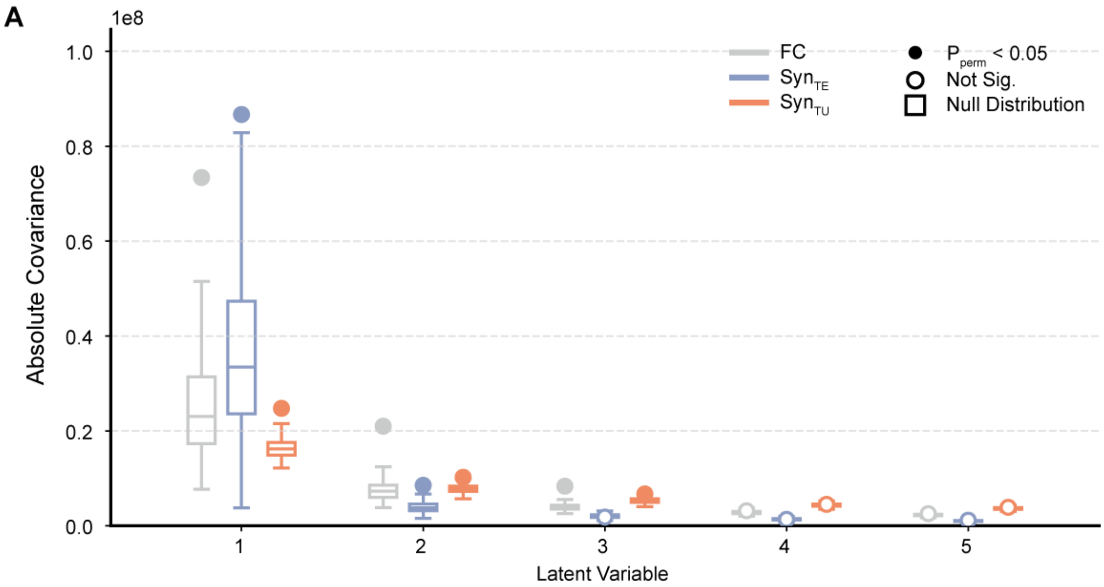


**Fig. S10 Effect sizes and statistical significance of latent variables across functional components.** The absolute covariance ($S^2$) is shown for the first five latent variables (LVs) derived from the PLS analysis linking brain connectivity networks to individual cognitive profiles. Results are systematically compared across empirical functional connectivity (FC), tract-explainable synchrony ($\mathrm{Syn}_{\mathrm{TE}}$), and tract-underexplained synchrony ($\mathrm{Syn}_{\mathrm{TU}}$). The corresponding empirical null distributions of $S^2$, generated via 1,000 permutation tests, are represented by boxplots for each LV and modality. Solid circles denote statistically significant LVs ($p_{\mathrm{perm}} < 0.05$), whereas open circles indicate non-significant effects.

#### Supplementary Tables

**Table S1 Mapping of HCP task variables to corresponding cognitive abilities.**

| HCP Task Variable | Cognitive Ability |
|---|---|
| PicVocab_Unadj | Vocabulary Comprehension |
| ReadEng_Unadj | Reading Decoding |
| PicSeq_Unadj | Episodic Memory |
| Flanker_Unadj | Inhibition |
| CardSort_Unadj | Cognitive Flexibility |
| ProcSpeed_Unadj | Processing Speed |
| PMAT24_A_CR | Fluid Intelligence |
| VSPLOT_TC | Spatial Orientation |
| IWRD_TOT | Word Memory |
| ListSort_Unadj | Working Memory |

#### References


[1] Van Essen, D. C. *et al.* The Human Connectome Project: A data acquisition perspective. *NeuroImage* **62**, 2222–2231 (2012).

[2] Hubbard, N. *et al.* Brain function and clinical characterization in the boston adolescent neuroimaging of depression and anxiety study. *NeuroImage: Clinical* **27**, 102240 (2020).

[3] Tian, X. *et al.* An integrated resource for functional and structural connectivity of the marmoset brain. *Nature Communications* **13**, 7416 (2022).

[4] Glasser, M. F. *et al.* The minimal preprocessing pipelines for the human connectome project. *NeuroImage* **80**, 105–124 (2013).

[5] Salimi-Khorshidi, G. *et al.* Automatic denoising of functional mri data: combining independent component analysis and hierarchical fusion of classifiers. *NeuroImage* **90**, 449–468 (2014).

[6] Tournier, J.-D. *et al.* Mrtrix3: A fast, flexible and open software framework for medical image processing and visualisation. *NeuroImage* **202**, 116137 (2019).

[7] Hawrylycz, M. J. *et al.* An anatomically comprehensive atlas of the adult human brain transcriptome. *Nature* **489**, 391–399 (2012).

[8] Arnatkeviciute, A., Fulcher, B. D. & Fornito, A. A practical guide to linking brain-wide gene expression and neuroimaging data. *NeuroImage* **189**, 353–367 (2019).

[9] Larivière, S. *et al.* The enigma toolbox: multiscale neural contextualization of multisite neuroimaging datasets. *Nature methods* **18**, 698–700 (2021).

[10] Hansen, J. Y. *et al.* Mapping neurotransmitter systems to the structural and functional organization of the human neocortex. *Nature Neuroscience* **25**, 1569–1581 (2022).

[11] Vaishnavi, S. N. *et al.* Regional aerobic glycolysis in the human brain. *Proceedings of the National Academy of Sciences of the United States of America* **107**, 17757–17762 (2010).

[12] Finnema, S. J. *et al.* Kinetic evaluation and test-retest reproducibility of [11C]UCB-J, a novel radioligand for positron emission tomography imaging of synaptic vesicle glycoprotein 2A in humans. *Journal of Cerebral Blood Flow and Metabolism: Official Journal of the International Society of Cerebral Blood Flow and Metabolism* **38**, 2041–2052 (2018).

[13] Zilles, K. & Palomero-Gallagher, N. Multiple transmitter receptors in regions and layers of the human cerebral cortex. *Frontiers in Neuroanatomy* **11**, 78 (2017).

[14] Goulas, A. *et al.* The natural axis of transmitter receptor distribution in the human cerebral cortex. *Proceedings of the National Academy of Sciences of the United States of America* **118**, e2020574118 (2021).

[15] Royer, J. *et al.* An open mri dataset for multiscale neuroscience. *Scientific Data* **9**, 569 (2022).

[16] Hill, J. *et al.* Similar patterns of cortical expansion during human development and evolution. *Proceedings of the National Academy of Sciences of the United States of America* **107**, 13135–13140 (2010).

[17] Shafiei, G. *et al.* Neurophysiological signatures of cortical micro-architecture. *Nature Communications* **14**, 6000 (2023).

[18] Tadel, F., Baillet, S., Mosher, J. C., Pantazis, D. & Leahy, R. M. Brainstorm: A user-friendly application for MEG/EEG analysis. *Computational Intelligence and Neuroscience* **2011**, 879716 (2011).

[19] Schaefer, A. *et al.* Local-Global Parcellation of the Human Cerebral Cortex from Intrinsic Functional Connectivity MRI. *Cerebral Cortex* **28**, 3095–3114 (2018).

[20] Fulcher, B. D. & Jones, N. S. *Hctsa*: A Computational Framework for Automated Time-Series Phenotyping Using Massive Feature Extraction. *Cell Systems* **5**, 527–531.e3 (2017).

[21] Markello, R. D. *et al.* Neuromaps: Structural and functional interpretation of brain maps. *Nature Methods* **19**, 1472–1479 (2022).

[22] Donoghue, T. *et al.* Parameterizing neural power spectra into periodic and aperiodic components. *Nature Neuroscience* **23**, 1655–1665 (2020).

[23] Pang, J. C. *et al.* Geometric constraints on human brain function. *Nature* **618**, 566–574 (2023).

[24] Wang, R. *et al.* Segregation, integration, and balance of large-scale resting brain networks configure different cognitive abilities. *Proceedings of the National Academy of Sciences of the United States of America* **118**, e2022288118 (2021).
